# Transient lung eosinophilia during breakthrough influenza infection in vaccinated mice is associated with protective and balanced Type 1/2 immune responses

**DOI:** 10.1101/2025.05.19.654943

**Authors:** Lauren A. Chang, Stephen T. Yeung, Prajakta Warang, Moataz Noureddine, Gagandeep Singh, Brett T. Webb, Michael Schotsaert

## Abstract

Eosinophils are agile cells that participate in a multitude of homeostatic and inflammatory responses in the lung, ranging from allergic asthma to antiviral defense against respiratory viral infection. In the context of vaccination followed by viral infection, such as breakthrough infection, eosinophils have been linked to aberrant Th2 responses like vaccine-enhanced respiratory disease (VAERD). Here, we demonstrate that the lung immune cell composition, cytokine and chemokine repertoire, histopathological profile, and systemic humoral response of breakthrough influenza infection in mice is distinct from that of primary influenza infection or allergic sensitization, canonical Type 1 and 2 immune responses, respectively. Longitudinal comparison of breakthrough infection with allergic sensitization and primary influenza infection demonstrated major differences in lung immunity between treatment groups in female, BALB/c mice. Breakthrough infection mice exhibit lung eosinophil infiltration that peaks at 7-10 days post-challenge, enriched for the Siglec-F^hi^ subset, but in the absence of overt pro-inflammatory cytokine/chemokine signals, high viral titers, severe lung lesions, goblet cell hyperplasia, allergic levels of total IgE, or enhanced morbidity. Multiparameter fluorescence imaging corroborated findings from flow cytometry and also unveiled interactions between CD101^+^Siglec-F^+^ cells with CD3^+^ cells in the lung tissue space. Imaging also revealed a marked absence of eosinophil or neutrophil extracellular traps in the lungs of breakthrough infection mice, in contrast with allergic sensitization and primary influenza infection, respectively. Altogether, our findings provide a deeper understanding of the kinetics and cell-cell interplay during non-pathological, balanced Type 1/2 immune responses in vaccinated hosts during breakthrough infection.

## Introduction

Humans are repeatedly infected with influenza over the course of their lifetime. Although seasonal vaccination is able to reduce the incidence of transmission and severe illness, current clinically approved vaccines have variable efficacy.^1–4^ Breakthrough infections of vaccinated individuals can still occur, but with substantially lower disease severity.^5^ This is partially conferred by immune cells like memory CD8 T cells, generated by previous vaccination or infection, which can kill infected cells by recognizing conserved epitopes presented in the context of major histocompatibility complex (MHC) proteins.^6^ Innate immune cells can also aid in control of virus during breakthrough infection, extrapolating from their roles during primary influenza infection.^7^ For example, neutrophils, monocytes, and alveolar macrophages (AMs) can assist by phagocytic clearance of infected dead cells from virus-infected lungs.^7,8^ Another innate immune cell, the eosinophil, also has many documented functions during primary influenza infection.^9–12^ Eosinophils can exert both direct antiviral functions, such as killing via release of granule proteins, or indirect functions such as priming T cells as non-professional antigen presenting cells.^13^ Eosinophils are also involved in the tissue repair response, such as after virus-induced damage.^14,15^

Despite the many antiviral capabilities of eosinophils, the appearance of these cells during respiratory viral breakthrough infection of vaccinated hosts has a negative connotation. This is in part due to the association of eosinophils with vaccine-enhanced respiratory disease (VAERD) following alum-adjuvanted formalin-inactivated respiratory syncytial virus vaccine (FI-RSV) trials in infants during the 1960s, where vaccinated children had severe, clinical disease following natural RSV infection, including more frequent hospitalization.^16,17^ VAERD is characterized by elevated levels of canonical Type 2 cytokines (IL-4, IL-5, IL-13), elevated eosinophil counts, and increased pathology in vaccinated populations after encountering wild-type virus relative to unvaccinated populations.^17^ All of these hallmarks, alongside increased mucin production and goblet cell hyperplasia, have been described in mouse models for RSV VAERD.^18^ However, thorough investigation has pinpointed Th2-biased CD4 T cell responses and generation of low-avidity non-neutralizing antibodies as major drivers of RSV VAERD etiology, and not eosinophils.^17–19^ Influenza VAERD has been demonstrated in pigs, although the likely cause appears to be elicitation of non-neutralizing antibodies rather than aberrant Th2-skewing of CD4 T cells.^20–22^

Our lab has optimized a breakthrough influenza infection model and have observed a dose-dependent recruitment of eosinophils to the lungs following challenge; mice that received three vaccinations had more lung eosinophils at 7 days post-challenge than mice that received one dose.^23^ Eosinophils did not appear to correlate with the severity of lesions, but rather with enhanced protection in stark contrast with VAERD. We further characterized this phenotype as a non-pathological enrichment of lung eosinophils, particularly that of the more inflammatory phenotype (Siglec-F^hi^ CD101^+^), in the absence of aberrant Type 1 or Type 2 inflammation, high concentrations of total serum IgE, or morbidity during influenza breakthrough infection.^24,25^ Despite the enrichment of eosinophils in the lung at 7 days post-challenge (DPC), no significant IL-5 or Eotaxin-1 (CCL11) signature was observed in lung homogenate supernatant cytokine/chemokine analysis, in contrast to observations in OVA-sensitized, allergic, positive control mice with robust eosinophilia which exhibited high concentrations of IL-5 and CCL11.^24^ Furthermore, the ratio of IgG2a/IgG1 as a surrogate of Th1/Th2 skewing did not track with the degree of eosinophil infiltration in breakthrough infection mice, since mice exhibiting the highest degree of lung eosinophilia had balanced IgG2a/IgG1 ratios rather than strongly IgG1-skewed ratios that would be indicative of Th2-biased immunity. Adjuvanting the vaccine with a strong Th1-skewing agent also did not deter eosinophil influx. Lung eosinophilia after breakthrough infection at 7 DPC was also observed in male BALB/c mice, indicating sex-independence. A vaccine-mismatched challenge in mice also recruited eosinophils to the lung. Collectively, our previous findings suggested that peripheral priming of hosts via intramuscular vaccination followed by a viral exposure in the lung resulted in non-allergic, non-pathogenic, complete resolution of respiratory viral disease, coinciding with eosinophilia rather than VAERD. The exact function of the recruited lung eosinophil subsets during breakthrough infection remains to be determined.

Given that previous studies by Choi et al.^23^ and Chang et al.^24^ assessed lung immunity at a singular time point (7 DPC), we conducted a longitudinal study with both the OVA allergic sensitization model and the influenza breakthrough infection model to better compare and contrast the kinetics of both immune responses. Primary influenza infection was also included as a condition to allow for direct comparison of breakthrough infection with a typical Type 1 immune reaction, alongside OVA allergic sensitization as a positive control for a canonical Type 2 response. After determining the time point where peak eosinophilia is observed, especially that of the Siglec-F^hi^ subset, we conducted multicolor fluorescence imaging. We visually confirmed eosinophilia in the lungs at 7 DPC, and observed granulocyte-T cell interactions as well. Furthermore, we saw neutrophil and eosinophil extracellular trap formation in the primary influenza infection and OVA-sensitized mice, respectively, but not in breakthrough infection mice. Therefore, eosinophil influx during influenza infection in the context of pre-existing immunity (breakthrough) may reflect different functionality compared to allergy-induced eosinophilia.

## Methods

### Study design

Female, 6-8 week old BALB/c mice were obtained from The Jackson Laboratory (Bar Harbor, ME). Mice were housed under specific pathogen-free conditions with food and water provided *ad libitum*. All experiments were approved by and performed according to the Icahn School of Medicine at Mount Sinai Institutional Animal Care and Use Committee. The vaccine used in this study was obtained through BEI Resources, NIAID, NIH: Fluzone® Influenza Virus Vaccine, 2005-2006 Formula, NR-10480. Mice were vaccinated intramuscularly in the hind limbs with a seasonal trivalent inactivated influenza virus vaccine (TIV; Fluzone® Influenza Virus Vaccine) containing an influenza A H1N1 component (A/New Caledonia/20/1999/IVR-116), influenza A H3N2 component (A/New York/55/2004/X-157 [an A/California/7/2004-like strain]), and influenza B component (B/Jiangsu/10/2003 [a B/Shanghai/361/2002-like strain]). PBS was injected as a negative control for vaccination. Non-adjuvanted TIV or PBS was administered intramuscularly to both hind limbs (50 µL/limb, 100 µL/mouse total). 5 mice were used per treatment group per time point.

### Influenza challenge

Mice were challenged with a sublethal (0.2 LD_50_) dose of H1N1 A/New Caledonia/20/1999 (NC99), as described previously for the breakthrough infection model.^23,24^ Briefly, mice were administered a mixture of ketamine and xylazine intraperitoneally (i.p.) for anesthesia. Mice then received a 50 µL intranasal (i.n.) challenge of either mouse-adapted egg-grown influenza virus or egg allantoic fluid as vehicle control. Body weights were recorded immediately before the challenge to define 100% body weight, then mice were weighed daily through 10 days post-infection to monitor morbidity. Additional body weight measurements were recorded for relevant groups at 28-30 days post-challenge prior to harvest.

### OVA sensitization

Mice were sensitized to OVA as described previously.^24^ Briefly, mice were sensitized twice with a 100 µL i.p. injection containing 20 µg of Imject™ ovalbumin (OVA) (Thermo Scientific, cat. no. 77120) adsorbed to alum, administered 1 week apart. Negative control mice received a 100 µL i.p. injection of PBS. One week after the final sensitization, mice were anesthetized as described above and i.n. challenged with 20 µg OVA diluted in PBS (final volume of 50 µL/mouse).

Control mice did not receive an intranasal challenge. Three mice were used per treatment group at each time point. Body weights were recorded as described above.

### Serum collection

Blood was collected from the submandibular vein and coagulated at 4°C overnight. The next day, coagulated blood was centrifuged at 400 x *g* for 5 minutes at 4°C. Serum was collected and stored at -20°C until analysis.

### Flow cytometry

Mice were euthanized with sodium pentobarbital. The left lung lobe was collected into a C-tube (Miltenyi Biotec, cat. no. 130-096-334) filled with 2.4 mL of 1X Buffer S from the Miltenyi Biotec Mouse Lung Dissociation Kit (cat. no. 130-095-927), diluted as per manufacturer’s instructions. 100 µL of Enzyme D and 15 µL of Enzyme A were spiked into each C-tube then digested on a Miltenyi Biotec gentleMACS Octo Dissociator with Heaters using the 37C_m_LDK_1 protocol. Digested lungs were then centrifuged at 400 x *g* for 5 minutes and supernatant was discarded. Pellets were resuspended in 5 mL of 1X Red Blood Cell (RBC) Lysis Buffer (8.29 g NH_4_Cl, 1.00 g KHCO_3_, and 200 µL of 0.5 M EDTA dissolved in ddH_2_O) then incubated at room temperature for 5 minutes, before centrifugation at 400 x *g* for 5 minutes. After RBC lysis, pellets were resuspended with 5 mL of PBS, passed through a 70 µm cell strainer (Greiner, cat. no. 542170), then centrifuged at 400 x *g* for 5 minutes. Supernatants were discarded and cell pellets were resuspended with 200 µL of Flow Cytometry (FC) Buffer (1% BSA, 2 mM EDTA in 1X PBS) before transferring the cell suspension to a 96-well V-bottom polypropylene plate (Thermo Scientific, cat. no. 249944). Plates were centrifuged at 400 x *g* for 5 minutes, supernatants were discarded, and pellets were stained for flow cytometry as described previously.^24^ Briefly, pellets were resuspended with 50 µL of Purified Rat Anti-Mouse CD16/CD32 Fc Block (clone 2.4G2, BD) diluted 1:100 in FC Buffer and incubated at room temperature for 5 minutes in the dark. Subsequently, 50 µL of surface staining antibody cocktail was added to each well of Fc blocked cells, bringing the total well volume to 100 µL. The cocktail contained the following antibodies and dyes: CD11c FITC (1:150, clone HL3, BD), CD125 PE (1:150, clone T21, BD), Siglec-F PE-CF594 (1:150, clone E50-2440, BD), Ly6G PerCP-Cy5.5 (1:150, clone 1A8, BD), CD101 PE-Cy7 (1:150, clone Moushi101, Invitrogen), CD11b APC (1:150, clone M1/70, BioLegend), MHC II Alexa Fluor 700 (1:150, clone M5/114.15.2, Invitrogen), CD62L APC-Cy7 (1:150, clone MEL-14, BD), and Fixable Viability Dye eFluor 450 (1:200, eBioscience). Cells were incubated with the surface staining cocktail at room temperature for 20 minutes in the dark. To wash, 120 µL of FC Buffer was added on top of the cells, followed by centrifugation at 400 x *g* for 5 minutes. Supernatants were decanted, plates were blotted on paper towels, then a second wash was performed by resuspending pellets with 200 µL of FC Buffer followed by centrifugation at 400 *x* g for 5 minutes. After washing, cell pellets were resuspended in 200 µL of FC Buffer and 5 µL of CountBright Absolute Counting Beads (ThermoFisher) were added to all samples. For single-stained compensation controls, 1 uL of antibody was added to 1 drop of UltraComp eBeads Plus Compensation Beads (ThermoFisher) for each antibody in the panel. Samples were acquired on a Beckman Coulter Gallios flow cytometer with Kaluza software. Data were analyzed using FlowJo v10 (BD) and raw data were compensated using AutoSpill. Following manual gating in FlowJo, data were visualized and statistically analyzed in Graphpad Prism version 10.2.1.

### Determination of infectious viral titer

Lung middle, inferior, and post-caval lobes were collected in 500 µL of sterile PBS in prefilled homogenizer bead tubes containing 3.0 mm high impact zirconium beads (Benchmark Scientific, cat. no. D1032-30). Samples were snap-frozen on dry ice on the day of harvest immediately after collection, then stored at -80°C until analysis. Samples were thawed on ice on the day of the assay, then homogenized and centrifuged at 10,000 x *g* for 5 minutes at 4°C to clarify lung homogenate supernatants.

To determine the 50% tissue culture infectious dose (TCID_50_), 2 x 10^4^ Madin-Darby canine kidney (MDCK) cells (ATCC, cat. no. CCL-34) were seeded per well at a final volume of 100 µL in Dulbecco’s Modified Eagle Medium (DMEM) (Gibco) containing 10% heat-inactivated fetal bovine serum (FBS) and 100 U/mL Penicillin-Streptomycin (Gibco) in a 96-well flat-bottom plate (Corning), then incubated at 37°C for 24 h. On the day of the assay, clarified lung homogenate supernatants were serially diluted 1:10 for a total of 8 dilutions in 96-well U-bottom plates (Corning) in serum-free DMEM containing 100 U/mL Penicillin-Streptomycin and 1 µg/mL TPCK trypsin: 8 µL of sample was diluted into 72 µL media for each dilution. After all samples were diluted, the serum-containing media was removed from the cell plates, and cell plates were then washed twice with 200 µL of sterile PBS per well then patted dry on paper towels. Immediately after washing, 50 µL of diluted sample was transferred from the dilution plates to the cell plates and incubated at 37°C for 1 h. After the incubation, 100 µL of serum-free media was added to each well of the cell plates. Plates were then incubated at 37°C for 72 h. After incubation, supernatants were discarded and cells were fixed with 200 µL of 5% formaldehyde in PBS per well and incubated at 4°C overnight. The next day, supernatants were discarded and plates were washed 3 times with 200 µL/well of PBS-T (1X PBS with 0.05% Tween-20). Plates were then blocked with 200 µL/well of 5% milk in PBS-T for 1 h at room temperature. After blocking, plates were washed 3 times as described and 100 µL/well of primary antibody solution (hyper-immune mouse serum diluted 1:500 in 5% milk in PBS-T) was added to the plates. After incubating for 1 h at room temperature, plates were washed 3 times as described. The goat anti-mouse IgG-HRP detection antibody (Abcam, cat. no. ab6823) was diluted 1:5000 in 5% milk in PBS-T, then 100 µL/well was added to the washed plates. After incubating for 1 h at room temperature, plates were washed 3 times as described. Plates were then washed 1 additional time with 1X PBS. After washing, 100 µL/well of 1-Step Turbo TMB Substrate (ThermoFisher, cat. no. 34022) was added to the plates and incubated for 20 minutes at room temperature in the dark. After the incubation, the reaction was quenched by adding 100 µL/well of ELISA Stop Solution (Invitrogen, cat. no. SS04). Plates were read on a BioTek Synergy Neo2 (Agilent) plate reader at 450 nm and 650 nm. Background-subtracted optical density values (OD_450-650nm_) were used for data analyses. The TCID_50_ was determined using the Reed-Muensch method.^26^ Positive wells were defined as wells containing an OD_450-650nm_ value >2X the average of the OD_450-650nm_ values of the media-only wells.

### Bead-based multiplex cytokine/chemokine assay

Lung lobes were collected and processed into clarified lung homogenate supernatants as described above for the TCID_50_ assay. Bead-based multiplex cytokine/chemokine assays were performed on the same day as the TCID_50_ assay to minimize sample freeze-thaw number. To measure a total of 27 lung cytokines and chemokines, we used the ProcartaPlex™ Mouse Cytokine & Chemokine Panel 1, 26plex kit (ThermoFisher, cat. no. EPX260-26088-901,) in conjunction with the IL-33 Mouse ProcartaPlex™ Simplex Kit (ThermoFisher, cat. no. EPX010-26025-901). Kits were combined and assay was performed as described by the manufacturer. For all incubation steps unless stated otherwise, plates were placed on an orbital shaker set to 300 rpm. Briefly, 100 µL/well of clarified lung homogenate supernatant was added to washed beads in a black, optical flat-bottom 96-well plate and incubated for 30 minutes at room temperature protected from light. After the room temperature incubation, plates were placed on a flat surface at 4°C for overnight incubation. The following day, the plate was incubated for 30 minutes at room temperature then washed 3 times with 150 µL/well of 1X Wash buffer, prepared according to the kit instructions. After washing, the 1X Detection Antibody cocktail was combined as recommended by the manufacturer and 25 µL/well was added. Plates were incubated at room temperature for 30 minutes, protected from light. After incubation, plates were washed 3 times as described and 50 µL/well of 1X Streptavidin-PE solution was added to the plates. Plates were incubated for 30 minutes at room temperature, protected from light. Plates were then washed 3 times as described and 120 µL/well of Reading Buffer was added. Plates were incubated for 5 minutes at room temperature and acquired on a Luminex 100/200 analyzer (Millipore) with xPONENT software (version 4.3). Extrapolated data were exported from the xPONENT software and visualized in GraphPad Prism version 10.2.1 or with *pheatmap* using the R computing language (ver. 4.05) in RStudio (ver. 1.4.1106).

### Tissues and histochemical staining

Right superior lobes were collected in 10% buffered formalin. Tissues were processed whole, embedded, sectioned at 4-5 microns and stained with hematoxylin and eosin (H&E) by routine methods. Alcian blue periodic acid Schiff (AB-PAS) stains were performed as previously described on 4-5 micron sections.^27^

### Histopathology

On H&E stained sections, the percent area affected, epithelial degeneration and necrosis, inflammation and atelectasis were graded on a 0-5+ scale as detailed in the below table. AB-PAS stained sections were graded on a 1-5+ scale based on the number of positively staining cells in airways using the following scale: 1+ = no positively cells, 2+ = occasional positively staining cells, 3+ = low numbers of positively staining cells, 4+ = moderate numbers of positively staining cells and 5+ = nearly confluent positively staining cells. Staining was categorized based on predominant cytoplasmic staining amongst the cells, either AB positive, PAS positive or mixed and presence or absence of a main bronchus within the section. One mouse was omitted from analyses of pathological data due to the presence of background pneumonia in the lobe, unrelated to treatment, as per veterinary pathologist’s recommendation.

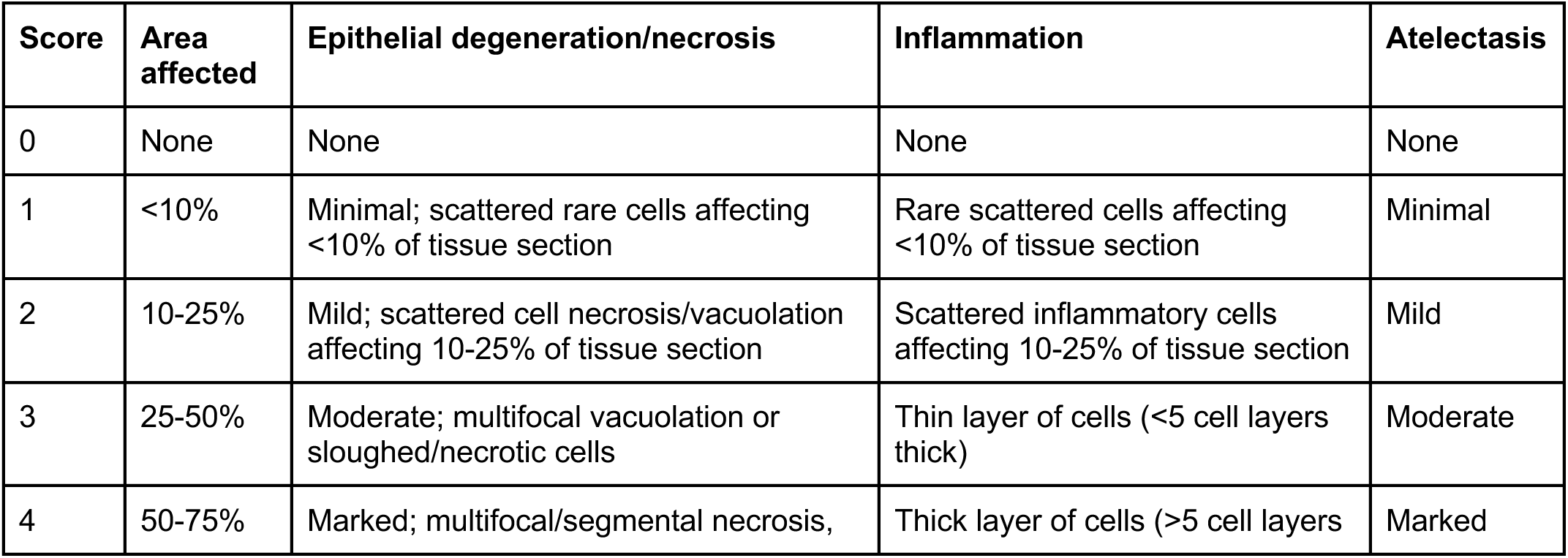
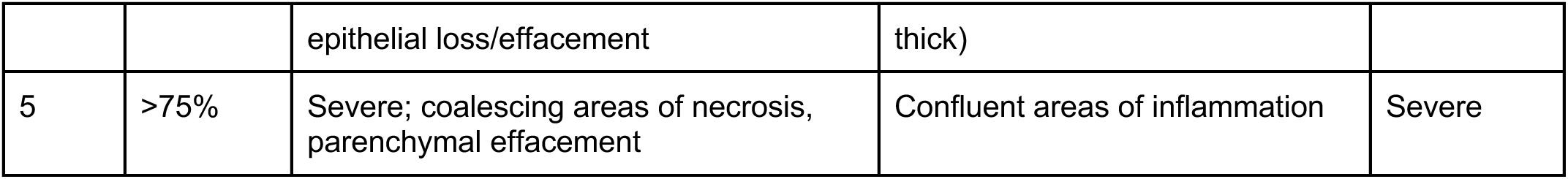

### Enzyme-linked immunosorbent assays (ELISA)

To measure OVA-specific antibody titers in the OVA-sensitized mice, Nunc MaxiSorp 96-well plates (ThermoFisher, cat.no. 456537) were coated with 100 µL/well of Imject™ OVA diluted to a concentration of 10 µg/mL in carbonate-bicarbonate buffer and incubated overnight at 4°C. The next day, plates were washed 3 times with 200 µL/well of PBS-T (1X PBS with 0.05% Tween-20) and patted dry on paper towels. Plates were then blocked with 200 µL/well of blocking buffer (5% milk in PBS-T). While plates were blocking for 1 hour at room temperature, sera were serially diluted in blocking buffer 4-fold a total of 7 times beginning from a 1:25 dilution (total IgG) or a 1:50 dilution (IgG1, IgG2a). Serial dilutions were performed in a 96-well V-bottom polypropylene plate (Thermo Scientific, cat. no. 249944). Plates were washed 3 times as described after blocking, then 50 µL/well of diluted sera was added to the plates, incubating for 1 h at room temperature. After incubation with serially diluted sera, plates were washed 3 times and 100 µL/well of diluted detection antibody was added: either goat anti-mouse IgG-HRP (1:5000, Abcam, cat. no. ab6823), goat anti-mouse IgG1-HRP (1:4000, SouthernBiotech, cat. no. 1071-05), or goat anti-mouse IgG2a-HRP (1:4000, SouthernBiotech, cat. no. 1081-05) diluted in blocking buffer. Plates were incubated for 1 hour at room temperature. After incubation with diluted detection antibody, plates were washed 3 times with PBS-T as described, followed by an additional wash with 1X PBS. Plates were then developed by adding 100 µL/well of 1-Step Turbo TMB Substrate (ThermoFisher), incubating for 20 minutes at room temperature in the dark. After incubation, 100 µL/well of ELISA Stop Solution (Invitrogen) was added to quench the reaction.

To measure vaccine-specific antibody titers in the breakthrough infection mice, Nunc MaxiSorp 96-well plates (ThermoFisher) were coated with 50 µL/well of TIV diluted 1:250 in carbonate-bicarbonate buffer and incubated overnight at 4°C. The next day, plates were washed and blocked as described above for 1 hour at room temperature. Sera were diluted in blocking buffer 5-fold a total of 7 times beginning from a 1:150 dilution (total IgG), or 3-fold a total of 7 times beginning from a 1:100 dilution (IgG1, IgG2a). As described above, plates were washed 3 times after blocking, then 100 100 µL/well of diluted sera was added to the plates, then incubated for 1 hr at room temperature. Plates were washed 3 times then 100 µL/well of diluted detection antibody was added as described above, for each respective isotype, and incubated for 1 hour at room temperature. After incubation with diluted detection antibody, plates were washed and developed as described above.

To measure total IgE, Nunc MaxiSorp 96-well plates (ThermoFisher) were coated with 100 µL/well of anti-mouse IgE antibody (Clone R35-72, BD) diluted to a concentration of 2 µg/mL in carbonate-bicarbonate buffer and incubated overnight at 4°C. The next day, plates were washed and blocked as described above for 1 hour at room temperature. As plates were blocking, each serum sample was diluted 1:50 in blocking buffer in technical duplicates in a 96-well V-bottom polypropylene plate (Thermo Scientific, cat. no. 249944). Unlabeled, purified mouse IgE (SouthernBiotech, cat. no. 0114-01) was diluted 4-fold a total of 11 times from a starting concentration of 4000 ng/mL to generate a standard curve. After blocking, plates were washed 3 times as described and 50 µL/well of diluted sera was added to plates and incubated for 1 h at room temperature. After incubation with sera, plates were washed and then incubated with 100 µL/well of goat anti-mouse IgE-HRP (SouthernBiotech, cat. no. 1110-05) diluted to 1:4000 in blocking buffer for 1 h at room temperature. After incubation with diluted detection antibody, plates were washed and developed as described above.

For all ELISA assays, plates were read on a BioTek Synergy Neo2 (Agilent) plate reader at 450 nm and 650 nm. Background-subtracted optical density values (OD_450-650nm_) were used for data analyses. Endpoint titers and interpolated total IgE concentrations were calculated in GraphPad Prism version 10.2.1.

### Principal component analysis

The following metrics from each mouse was integrated into a singular data frame for principal component analysis (PCA): flow cytometry absolute numbers (5 populations of interest); body weight percentages (up to 12 timepoints); concentrations of 27 different cytokines and chemokines measured from clarified lung homogenate supernatants; histopathology scores for 19 different metrics, including total score; goblet cell scores; and total IgE concentration (3 timepoints) for a total of 67 different metrics. OVA- and vaccine-specific total IgG, as well as TCID_50_ values were omitted from analyses to facilitate concomitant antigen-agnostic comparison of host immune profiles observed in the OVA sensitization and breakthrough infection studies. Data visualization and analysis were conducted using *prcomp* in the R computing language (ver. 4.05) in RStudio (ver. 1.4.1106).

### Tissue preparation for immunofluorescence and confocal microscopy

Briefly, tissues were fixed in paraformaldehyde, lysine, and sodium periodate buffer (PLP, 0.05 M phosphate buffer, 0.1 M L-lysine, pH 7.4, 2 mg/mL NaIO4, and 10 mg/mL paraformaldehyde) overnight at 4°C followed by 30% sucrose overnight at 4°C and subsequent embedding in OCT media. 20 µm frozen tissue sections were sectioned using a Leica CM3050S cryostat (Leica Biosystems Inc). FcR blocked with anti-CD16/32 Fc block antibody (clone 93, BioLegend) diluted in 1X PBS containing 2% donkey serum and 2% FBS for 1 h at room temperature. Sections were stained with Siglec-F BV421 (Clone E50-2440, BD), CD11b AF488 (Clone M1/70, BioLegend), CD11c PE (Clone N418, Invitrogen), Ly6G APC (Clone 1A8, BioLegend), Ly6G BV510 (Clone 1A8, BD), CD62L AF488 (Clone MEL-14, Southern Biotech), CD101 PE (Clone Moushi101, Invitrogen), CD11c eFluor 615 (Clone N418, Invitrogen), CD125 APC (Clone DIH37, BioLegend), CD3e APC (Clone 145-2C11, BioLegend), B220 V500 (Clone RA3-6B2, BD), Ly6G AF488 (Clone 1A8, BioLegend), anti-EPX (Clone MM25.8.2.2, antibody kindly provided by Dr. Elizabeth Jacobsen), or anti-Citrullinated Histone H3 (Abcam) diluted in PBS containing 2% donkey serum, 2% FBS, and 0.5% Fc block for 1 h at room temperature. Sections were subsequently washed with 1X PBS to remove unbound antibody. Following, sections were stained with goat anti-mouse AF488 (Invitrogen) and Goat anti-rabbit AF546 (Invitrogen) secondary antibodies diluted in 1X PBS containing 2% donkey serum, 2% FBS, and 0.5% Fc block for 1 h at room temperature. Sections were nuclear stained with DRAQ7 (ThermoFisher, Waltham, MA, USA) for 5 minutes at room temperature and then washed with 1X PBS. Slides were subsequently washed with 1X PBS and cover slipped using Immunmount mounting medium (Fisher Scientific) and Cover Glasses with a 0.13 to 0.17 mm thickness (Fisher Scientific). Fluorescence was detected with a Zeiss LSM 880 confocal microscope (Carl Zeiss) equipped with 405, 488, 514, 561, 594, and 633 nm solid-state laser lines, a 32-channel spectral detector (409–695 nm), and 10×0.3, 20x Plan-Apochromat 0.8, 40x, and 6331.40 objectives. Zen Black (Carl Zeiss) software suite was used for data collection. The imaging data were processed using Imaris software version 8.3.1 (Bitplane USA; Oxford Instruments). Image analysis by FIJI ImageJ and QuPath were as previously described.^28,29^ Following Imaris processing, images were analyzed using FIJI ImageJ version 2.16.0/1.54p (NIH, Bethesda, MD, USA) for number of immunostained colocalization and cell-cell interactions by using FIJI Cell Counter plugin. Image analysis was conducted on acquired images as follows: two sections per slide per animal of *n*=2-3 animals.

## Results

### Kinetics and enrichment dynamics of multiple key immune cell populations in the lung differ drastically during allergic sensitization compared to breakthrough influenza infection

We have established a breakthrough influenza infection model using the 2005-2006 trivalent inactivated influenza vaccine (TIV; Fluzone) as the model vaccine: mice are vaccinated with TIV, then intranasally challenged with a sublethal (0.2 LD_50_) dose of a vaccine-matched influenza virus, H1N1 A/New Caledonia/20/1999 (NC99). Previously, we have shown that replicating virus is observed early after challenge at 2-3 DPC, but rapidly cleared and undetectable by 7 DPC in vaccinated hosts, in stark contrast to mice without prior vaccination undergoing primary infection which do not clear the infection by 7 DPC and still have detectable viral titers.^23,24^

We previously observed a 2- to 4-fold increase in lung eosinophil numbers in mice experiencing breakthrough infection, with a marked enrichment in the Siglec-F^hi^ subset.^24^ This lung eosinophilia was not accompanied by an overt Th2 cytokine module, substantial host weight loss, abrogation of alveolar macrophage (AM) numbers, or high concentrations of total serum IgE, unlike what is observed in typical, pathological Type 2 responses such as allergy or vaccine enhanced respiratory disease (VAERD). This non-pathological lung eosinophilia was also present irrespective of adjuvant: inclusion of a Th2-skewing agent such as alum or a potent Th1-skewing agent such as an amphiphilic conjugate of the imidazoquinoline TLR7/8 agonist connected to poly(ethylene glycol) and cholesterol (IMDQ) during vaccination elicited similar numbers of lung eosinophils upon breakthrough infection. This eosinophil recruitment also appeared to be sex-independent, as it was observed in male mice as well.^24^

Thus far, all characterization of immunity has been conducted at 7 days post-challenge (DPC). We expanded our analyses to include additional time points to pinpoint when peak eosinophilia occurs during the acute phase of inflammation, and to observe if there are any clear cytokine and chemokine recruitment signals. We also included a later harvest time point at 28 DPC to evaluate if the inflammatory phenotypic shift of the lung eosinophil population is more long-lived than the reported half-life of 1.5-8 days in the lung, which could point to prolonged recruitment, retention, or local renewal.^14,30,31^ To better understand differences in lung immune cell kinetics, we compared and contrasted a typical, pathological Type 2 response model, ovalbumin (OVA) sensitization, with our experimental breakthrough influenza infection model at days 1, 3, 7, 10, and 28 DPC (Fig. 1A-B). Using flow cytometry, we quantified multiple key immune cell populations in the lung: neutrophils, alveolar macrophages, and eosinophils. Gating strategy is as described in Fig. S1.

**Figure 1.**
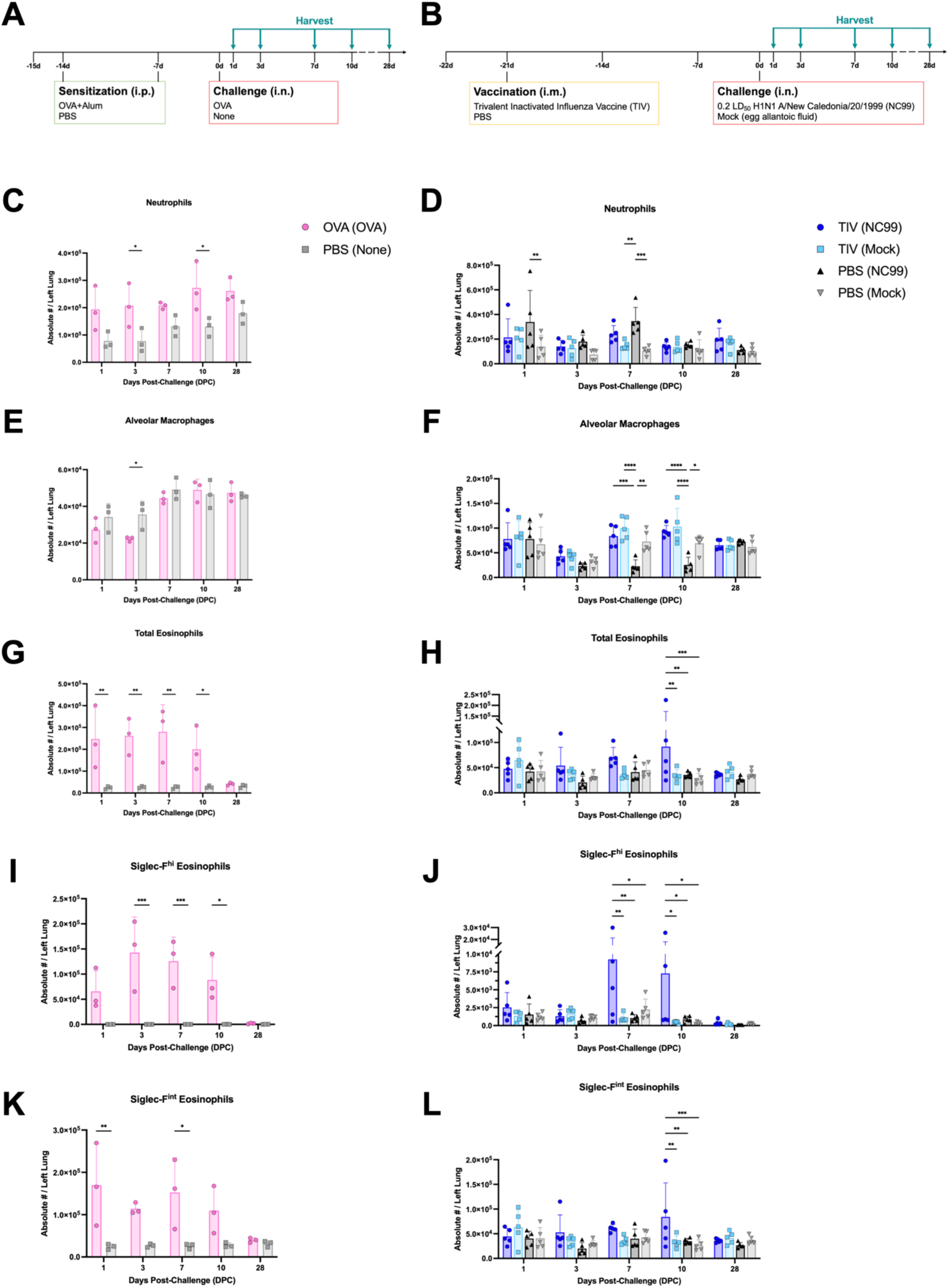
OVA sensitization and breakthrough influenza infection exhibit different immune cell kinetics during acute lung inflammation. Study design for (**A**) OVA sensitization and (**B**) breakthrough infection longitudinal studies. Absolute numbers of (**C, D**) neutrophils (Ly6G^+^), (**E, F**) alveolar macrophages (Ly6G^-^ CD11b^-^ CD11c^+^ Siglec-F^+^ MHC II^hi^), (**G, H**) total eosinophils (Ly6G^-^ CD11b^+^ CD11c^-^ CD125^int^ Siglec-F^+^), (**I, J**) Siglec-F^hi^ eosinophils, and (**K, L**) Siglec-F^int^ eosinophils quantified from the (**C, E, G, I, K**) OVA sensitization or (**D, F, H, J, L**) breakthrough infection longitudinal studies. Bar plots and error bars represent mean ± SD, with each symbol representing an individual mouse. Statistical significance was determined via ordinary two-way ANOVA (**C, E, G, I, K**) with Šídák’s multiple comparisons test with a single pooled variance or (**D, F, H, J, L**) with Tukey’s multiple comparisons test with a single pooled variance. ****P < 0.0001, ***P = 0.0001 to 0.001, **P = 0.001 to 0.01, *P = 0.01 to 0.05.

Neutrophils are key innate immune cells that can rapidly respond to inflammatory insults. Neutrophil recruitment to the lungs has been implicated in both allergic asthma and respiratory viral infections, with differing effects on host outcome.^32,33^ Given their importance in the lung immune response, we quantified total neutrophil counts in both the OVA sensitization model and breakthrough infection model (Fig. 1C-D). In OVA sensitized mice, a 1.5- to 3.2-fold enrichment in neutrophil count was observed through 10 DPC compared to the PBS control group that received no intranasal challenge, with significant differences at 3 and 10 DPC (Fig. 1C). Lung neutrophil numbers were most elevated at 10 DPC in the OVA sensitized group and sustained, even at 28 DPC, compared to PBS control group which had steadily increasing neutrophil counts through the end of the study. Mice that received PBS vaccination and were challenged with NC99 H1N1 virus, essentially undergoing a primary influenza infection, had significantly higher counts of lung neutrophils at 1 and 7 DPC compared to the PBS-vaccinated mock-challenged controls (Fig. 1D). TIV-vaccinated NC99-challenged group, which modeled breakthrough infection, had relatively stable numbers of lung neutrophils throughout the course of the study, similar to the TIV-vaccinated mock-challenged group and PBS-vaccinated mock-challenged group. Within the two TIV-vaccinated groups, neutrophil counts were not significantly different, irrespective of if the mice were challenged or not.

Next, we quantified the number of AMs in the lungs of mice undergoing OVA sensitization or breakthrough infection (Fig. 1E-F). AMs are tissue-resident macrophages in the alveolar space and have a multitude of homeostatic and inflammatory functions. AMs serve as the first line of defense against perturbations in the respiratory tract and are nimble cells, capable of promoting both pro- and anti-inflammatory states in allergic asthma, respiratory viral infections, and more.^34,35^ In OVA-sensitized mice, the number of lung AMs was significantly reduced at 3 DPC compared to PBS control mice, although counts rapidly recovered to the level of control mice by 10 DPC (Fig. 1E). This is in line with findings demonstrating that AMs can die during airway allergic responses.^36^ In the breakthrough infection model, both the NC99- and mock-challenged groups, irrespective of vaccine regimen, had lower numbers of AMs in the lung at 3 DPC relative to 1 DPC (Fig. 1F). However, for all groups except for PBS-vaccinated NC99-challenged primary influenza infection mice, AM numbers were similar to that of the unchallenged PBS negative control in the OVA sensitization model, suggesting that the counts at 3 DPC, while lower than 1 DPC, are still within the homeostatic range or were transiently reduced as a result of the intranasal instillation of challenge material (Fig. 1E). AM count rapidly increased by 7 DPC for all groups in the breakthrough infection model, except for the PBS-vaccinated NC99-challenged primary infection mice, which had a highly significant reduction in AM numbers through 10 DPC. In contrast, TIV-vaccinated NC99-challenged breakthrough infection mice had no such reduction in AMs, with numbers resembling that of the TIV-vaccinated mock-challenged healthy controls at all time points in the study. This corroborates previous studies, where we and others have demonstrated that vaccine-mediated immunity prevents AM death upon subsequent influenza infection.^23,24,37–40^ By 28 DPC, all groups had equivalent numbers of AMs.

Eosinophils are recruited during Type 2 responses, such as allergy and asthma, where eosinophil degranulation or release of cytokines can facilitate immunopathologic disease.^30,41,42^ Lung eosinophil influx has also been associated with respiratory viral breakthrough infection, particularly in the case of RSV VAERD, although they are not directly responsible for the enhanced disease phenotype; Th2 CD4 T cells are the main drivers of RSV VAERD.^17–19^ Eosinophils during influenza breakthrough infection have been investigated by our group, and in contrast with observations in the RSV model and allergy models, lung eosinophils do not correlate with enhanced disease, but rather with protection.^23,24^ As such, we were interested in quantifying the number of lung eosinophils following OVA allergic sensitization and comparing it with the breakthrough infection model, to understand the differences in kinetics, magnitude of recruitment, and phenotype (Fig. 1G-H).

After OVA challenge, all OVA-sensitized mice had a substantial enrichment in lung eosinophil count compared to the PBS control group during the acute phase of inflammation (Fig. 1G). OVA-sensitized mice had 9.6- to 10.7-fold more eosinophils than the PBS control at 1-7 DPC. By 10 DPC the number of eosinophils was lower than in earlier time points, although there were still 7.1-fold more eosinophils in the OVA sensitized group than the PBS controls. By 28 DPC, the eosinophil numbers were not significantly different between the two groups. For the breakthrough infection model, the TIV-vaccinated NC99-challenged breakthrough infection group generally had the highest numbers of lung eosinophils compared to other groups during the acute stage of infection (Fig. 1H). Eosinophil counts were 2.6-fold higher in the breakthrough infection group compared to the primary infection group at 3 DPC. There was a 1.8-fold enrichment in eosinophils in the breakthrough infection group that received a sublethal NC99 challenge, compared to the mock-challenged controls at 7 DPC. This is in line with previous experiments we conducted at 7 DPC.^24^ The breakthrough infection group also had a statistically significant, 1.6- to 2.2-fold greater enrichment in eosinophil numbers at 10 DPC relative to other groups. By 28 DPC, all four treatment groups had similar numbers of eosinophils. In terms of kinetics, only the breakthrough infection group had a distinct rise-fall in eosinophil numbers, peaking at 10 DPC, while all other groups were relatively stable in eosinophil counts throughout the entire study.

Within the lung eosinophil population, we quantified two subpopulations: Siglec-F^hi^ eosinophils and Siglec-F^int^ eosinophils. Siglec-F^hi^ eosinophils are thought to be the more activated, inflammatory subset while Siglec-F^int^ eosinophils have been described to be the more resting, homeostatic phenotype in the lungs.^25^ CD101 expression has also been used to subset eosinophils, with the CD101^+^ eosinophils corresponding to more pro-inflammatory functions and prevalent in pathological Type 2 responses, such as allergy and helminth infections.^25^ CD101^+^ eosinophils have also been observed during breakthrough infection by our group, although they did not appear to correlate with host pathology.^24^ We observed that 75-90% of the Siglec-F^hi^ eosinophils were CD101^+^ at any given time point, in alignment with literature (Fig. S2A-B).^25,43–45^ In the OVA-sensitized mice, Siglec-F^hi^ eosinophils were robustly recruited to the lungs 1-10 DPC, whereas this subset of eosinophils did not have a substantial presence in PBS control mice that received no intranasal challenge throughout the entire study (Fig. 1I). Numbers of this subset peaked in the OVA-sensitized mice at 3 DPC, and gradually began to fall through 10 DPC. Although Siglec-F^hi^ eosinophils numbers were largely reduced by 28 DPC, OVA-sensitized mice still had 9.8-fold higher Siglec-F^hi^ eosinophils in the lung compared to the PBS control mice. Siglec-F^hi^ eosinophils were also the majority of the total eosinophil population 3-10 DPC in the OVA-sensitized mice, indicative of a phenotypic shift towards or preference for recruitment of this subset during acute allergic inflammation (Fig. S3A). For the breakthrough infection model, we observed all groups had some degree of recruitment of Siglec-F^hi^ eosinophils to the lung after any intranasal challenge (Fig. 1J). This subset was most prevalent in the breakthrough infection group alone, with significantly higher numbers than all other treatment groups at 7 and 10 DPC. Similar to kinetics seen in the OVA-sensitization study, the number of Siglec-F^hi^ eosinophils was much lower in all groups by 28 DPC compared to earlier time points in the study. Of note, the number of Siglec-F^hi^ eosinophils was >2-fold higher in the TIV-vaccinated NC99-challenged breakthrough infection mice compared to all other groups at 28 DPC, even at this later time point. Although Siglec-F^hi^ eosinophils were never the majority of the total eosinophil population for any of the treatment groups in the breakthrough infection model, this subset was the significantly enriched for at 7 DPC in the breakthrough infection group only: Siglec-F^hi^ eosinophils were 10.8% of eosinophils for TIV-vaccinated NC99-challenged mice compared to the TIV-vaccinated mock-challenged (2.9%), PBS-vaccinated NC99-challenged (4.7%), and PBS-vaccinated mock-challenged (2.7%) groups (Fig. S3B). Mirroring observations in Siglec-F^hi^ eosinophil counts, the frequency of this subset remained elevated in only the breakthrough infection group (1.1%) at 28 DPC, while <1% of total eosinophils were Siglec-F^hi^ eosinophils in all other groups by this time point.

Next, we quantified the number of Siglec-F^int^ eosinophils in the lungs for both models (Fig. 1K-L). In the OVA-sensitization model, numbers of Siglec-F^int^ eosinophils were elevated from 1-10 DPC and were the highest on 1 and 7 DPC (Fig. 1K). In contrast, PBS control mice had relatively stable numbers of Siglec-F^int^ eosinophils the entire study. Unlike what was seen for the Siglec-F^hi^ eosinophils, where numbers were still elevated above the control group at 28 DPC, the numbers of Siglec-F^int^ eosinophils were similar to that of the controls by this later time point. Although there was also a dramatic increase in the numbers of this subset, Siglec-F^int^ eosinophils represented <50% of total eosinophils at 3-10 DPC due to the increase in Siglec-F^hi^ eosinophils in the OVA-sensitized mice. Otherwise, in PBS control mice that received no intranasal challenge, >99% of eosinophils have the Siglec-F^int^ phenotype (Fig. S3C). In the breakthrough infection model, Siglec-F^int^ kinetics for each treatment group largely resembled that of the total eosinophil population (Fig. 1H, L). The enrichment in Siglec-F^int^ eosinophils was most apparent at 7 and 10 DPC: the TIV-vaccinated NC99-challenged breakthrough infection group had, respectively, 1.7- or 2.5-fold higher counts of Siglec-F^int^ eosinophils than the TIV-vaccinated mock-challenged control group. Breakthrough infection mice had significantly greater numbers of this subset than all other groups at 10 DPC. By 28 DPC, all groups had similar numbers of Siglec-F^int^ eosinophils, unlike what was observed for the Siglec-F^hi^ subset. We saw that Siglec-F^int^ eosinophils represented less of the total eosinophils at 7 DPC, corresponding to the significantly increased frequency of Siglec-F^hi^ eosinophils (Fig. S3D).

In summary, drastically different immune cell population dynamics were observed for both models used in this study. OVA-sensitization presented with the hallmarks of pathogenic Type 2 immune responses: exuberant lung eosinophilia for both subsets, transient loss of AMs, and enrichment in neutrophilic inflammation as well. In contrast, primary influenza infection in hosts without pre-existing vaccine immunity exhibited neutrophilia, marked and sustained loss of AMs, but no eosinophilia. Breakthrough infection of vaccinated hosts following sublethal viral challenge presented with a lung cellular composition that had elements of both the OVA-sensitized, Type 2 response and the primary influenza infection Type 1 response: mild neutrophil enrichment was observed, relative to vaccinated and mock-challenged controls, but not to the same extent as primary influenza infection mice; no dramatic loss of AMs was observed; and total eosinophil counts were significantly higher in this group at 7-10 DPC, with a substantial enrichment for the Siglec-F^hi^ subset, although not to the same extent as the OVA-sensitized mice.

### OVA-sensitized allergic mice and unvaccinated influenza infected mice have exuberant pro-inflammatory cytokine and chemokine expression in the lung during the acute phase of inflammation

As a snapshot of the inflammatory profile of the lungs, we next measured the concentrations of 27 different cytokines and chemokines in lung homogenate supernatants for both models. In the OVA sensitization study mice, we saw a very distinct pro-inflammatory cytokine and chemokine profile most prevalent at 1 DPC in the OVA-sensitized mice, which was not observed in the PBS control mice that did not receive an intranasal challenge (Fig. 2A, S4). OVA-sensitized mice had high concentrations of IL-1β, IL-6, IL-18, IL-27, GM-CSF, CCL2, CCL3, CCL4, CXCL1, and CXCL2. Type 1 cytokines such as TNF-α and IL-12, but not IFN-γ, were also substantially elevated in OVA-sensitized mice at 1 DPC but not in PBS control mice. Canonical Type 2 cytokines such as IL-4, IL-5, and IL-13 were significantly higher at 1 DPC in the OVA-sensitized mice and remained higher than concentrations observed in the PBS control mice through 7 DPC. Interestingly, CCL11 (Eotaxin-1) did not appear to have a strong signal at the acute time points when peak lung eosinophilia was observed in OVA-sensitized mice.

**Figure 2.**
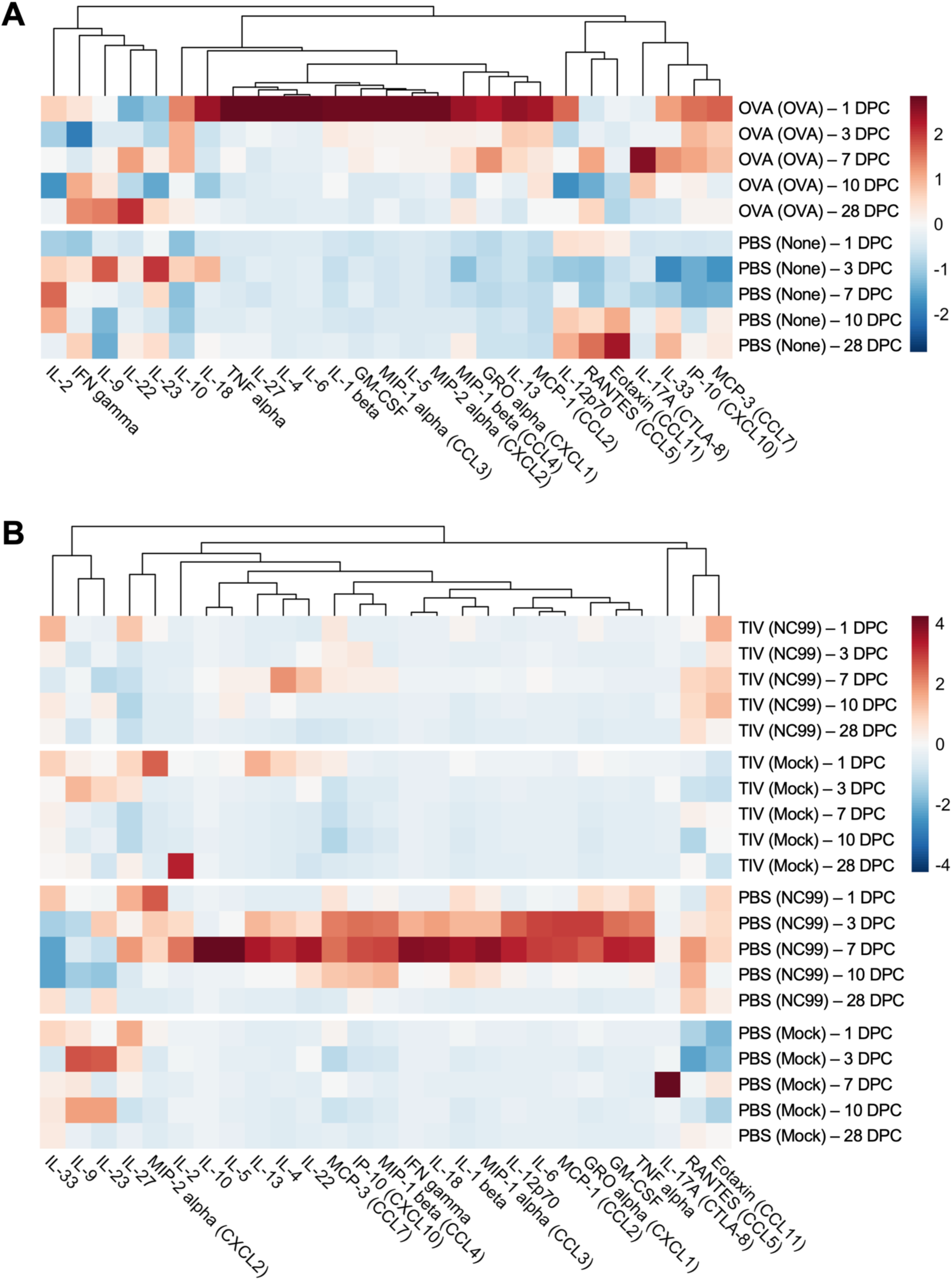
Breakthrough infection does not exhibit a distinctly Th1 or Th2 lung cytokine and chemokine module during acute and late infection, unlike OVA-sensitization and primary influenza infection. Heat map of net mean fluorescence intensity (MFI) of each analyte, *z*-scored by column (**A**) OVA sensitized mice and controls and (**B**) breakthrough infection mice and controls. Each square represents the average net MFI for the treatment group at that time point (*n* = 3-5).

Distinct cytokine and chemokine profiles were observed in the breakthrough infection model, contrasting with observations in the OVA-sensitized mice (Fig. 2B, S5). Many pro-inflammatory cytokines and chemokines were highly expressed in the PBS-vaccinated NC99-challenged primary infection mice at 3 and 7 DPC at significantly higher concentrations than all other groups: IL-1β, IL-6, IL-18, IL-22, IL-27, CCL2, CCL3, CXCL2, CCL4, CCL5, CCL7, CXCL1, and CXCL10. Canonical Type 1-associated cytokines such as IFN-γ, TNF-α, IL-2 and IL-12, were most prominent in this group. This is in line with expectations since this group was undergoing a primary viral infection. Additionally, Type 2 cytokines such as IL-4 and IL-13 were also present in the primary infection mice at these two time points, and with significantly higher IL-5 expression at 7 DPC than all other groups. Anti-inflammatory cytokine IL-10 was most highly expressed at 7 DPC in primary infection mice. Concentrations of the alarmin IL-33 were depressed in the primary infection mice most prevalently at days 3-10 post-challenge, ultimately recovering to the levels of all other treatment groups by 28 DPC. In stark contrast, the TIV-vaccinated NC99-challenged breakthrough infection mice had a substantially less inflammatory cytokine and chemokine profile in the lungs compared to primary infection mice. Type 1 cytokines were slightly elevated in TIV-vaccinated NC99-challenged breakthrough infection mice compared to TIV-vaccinated mock-challenged controls, but still significantly lower when compared to the PBS-vaccinated NC99-challenged primary infection mice. Similarly, low levels of Type 2 cytokines were observed in the breakthrough infection mice, but concentrations were still significantly higher in the primary infection mice. Concentrations of IL-4 were higher in the breakthrough infection mice compared to the mock-challenged controls at 7 DPC, but still significantly lower than the primary infection mice. CCL11 concentrations were significantly elevated in the breakthrough infection mice at 1 DPC compared to the primary infection mice, but otherwise were at similar concentrations as all other treatment groups at the remainder of the time points. Of note, peak CCL11 concentrations preceded the peak lung eosinophilia, observed at 7-10 DPC, by several days in the breakthrough infection mice. IL-27 concentrations in the breakthrough infection mice were significantly lower than that of PBS-vaccinated mock-challenged controls at 3 DPC, and significantly lower than that of TIV-vaccinated mock-challenged controls at 28 DPC.

Complementing our cellular quantification, we observed a distinct Type 2 cytokine and chemokine module for OVA-sensitized mice, a clear Type 1 profile for primary infection mice, and a more balanced, muted inflammatory profile for the breakthrough infection mice. Peak cytokine and chemokine signatures were observed at more acute time points following intranasal challenge, such as 1 DPC for the OVA-sensitized mice and 3-7 DPC for the primary infection mice.

### Host morbidity corresponds with peak lung inflammatory profile for OVA-sensitized mice, and viral titers for breakthrough and primary influenza infection mice

To bolster our understanding of general host health status beyond lung immunity, we used body weight as a metric of morbidity for both models. Despite exuberant recruitment of eosinophils to the lung, compounded by neutrophilia and transient loss of alveolar macrophages, OVA-sensitized mice did not have drastic reductions in body weight, unlike what is observed during severe respiratory viral infection (Fig. 3A). There was statistically significant weight loss on 1-2 DPC, with OVA-sensitized mice losing <3% body weight while PBS control mice did not. However, this difference may be due to sedation since PBS control mice received no intranasal challenge and therefore were not sedated. At 28 DPC, OVA-challenged mice had significantly higher body weights than the PBS control mice.

**Figure 3.**
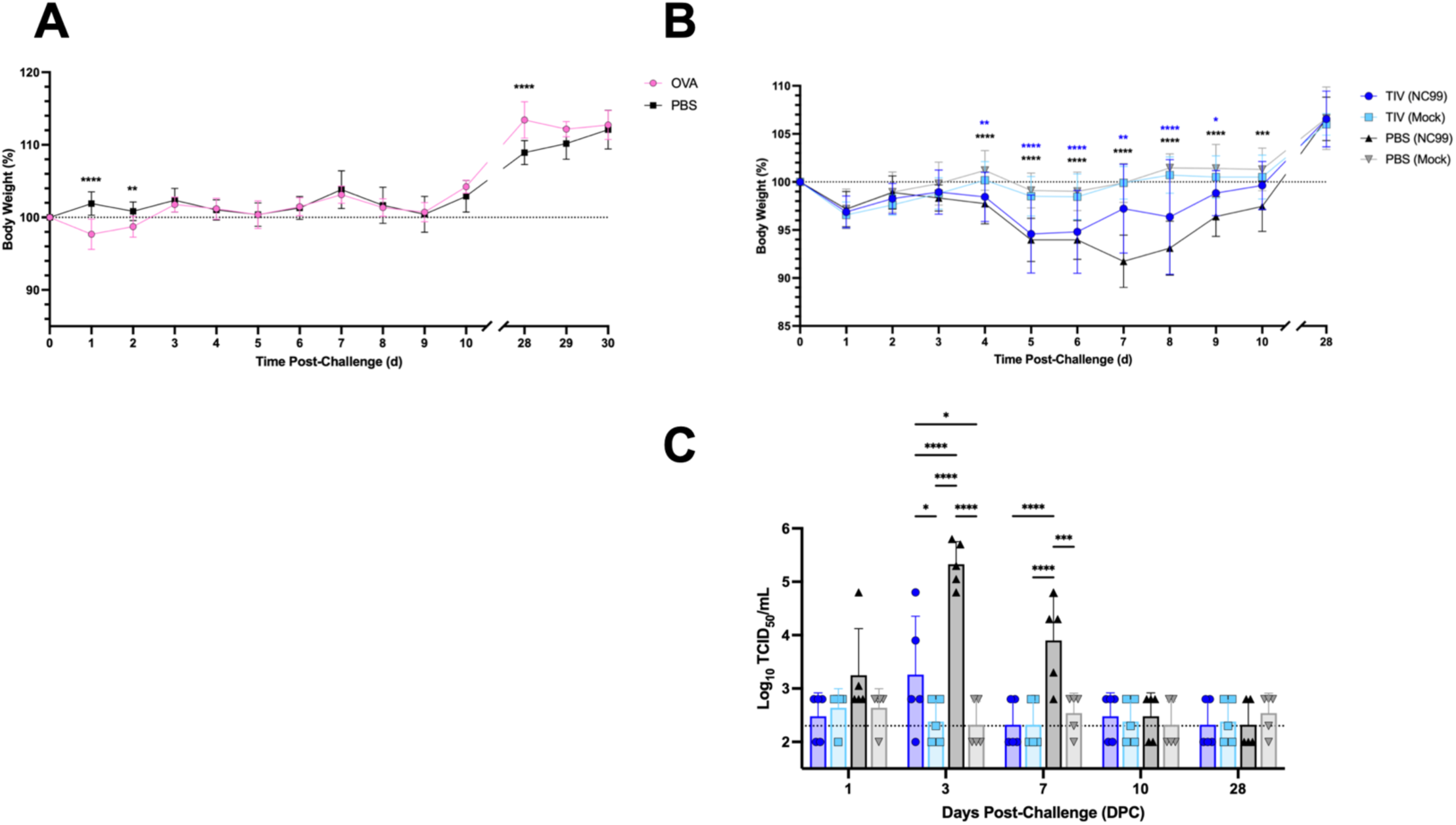
Host body weight does not correlate to lung immune inflammation in OVA-sensitized mice, but does reflect severity of viral load in mice experiencing breakthrough or primary influenza infection. Morbidity for (**A**) OVA sensitized mice and controls or (**B**) breakthrough infection mice and controls. (**C**) Replicating viral load in the lungs was determined via TCID_50_ for breakthrough infection mice and controls. Dotted lines denote (**A**, **B**) 100% body weight or (**C**) LOD of 200 TCID_50_/mL. Bar plots and error bars represent mean ± SD, with each symbol representing an individual mouse. Statistical significance was determined via ordinary two-way ANOVA with (**A**) Šídák’s multiple comparisons test with a single pooled variance, (**B**) Dunnett’s multiple comparisons test with a single pooled variance using the PBS (Mock) group as the comparator for all other groups, or (**C**) Tukey’s multiple comparisons test, with a single pooled variance. Asterisks are color-coded by group in (**B**). ****P < 0.0001, ***P = 0.0001 to 0.001, **P = 0.001 to 0.01, *P = 0.01 to 0.05.

For the breakthrough infection study, both the TIV- and PBS-vaccinated groups that received the sublethal NC99 challenge lost weight (Fig. 3B). PBS-vaccinated NC99-challenged primary infection mice continually lost weight through 7 DPC before recovering, with a maximum loss of 8.3% body weight at 7 DPC. In contrast, TIV-vaccinated NC99-challenged breakthrough infection mice began to recover at 5 DPC and had a maximum weight loss of 5.4% at this time point. By 10 DPC, breakthrough infection mice had similar weights as the PBS-vaccinated mock-challenged control group, whereas primary infection mice still had significantly lower body weights. By 28 DPC, both virus-challenged groups had equivalent body weights as the mock-challenged control groups. Our morbidity data was corroborated by our viral load measurements. We found that primary infection mice had the highest viral burden at 3 DPC, significantly higher than all other groups, including mice with vaccine protection (Fig. 3C). There was a low level of replicating virus measured in the lungs of TIV-vaccinated NC99-challenged mice at 3 DPC, confirming breakthrough infection. By 7 DPC, breakthrough infection mice had TCID_50_ measurements equivalent to the mock-challenged groups, indicating rapid control of the virus. In contrast, primary infection mice that received PBS-vaccination and had no prior immunity against influenza still had significantly higher viral load at 7 DPC when compared to all other groups. By 10 DPC, primary infection mice had TCID_50_ measurements similar to the TIV-vaccinated NC99-challenged group as well as the mock-challenged groups, indicating control of viral replication by this time point.

### Lung disease is most severe for OVA-sensitized mice and primary influenza infection mice at acute time points, while breakthrough infection mice are protected from severe disease and do not exhibit extensive mucin staining

After quantifying lung immune cell populations, inflammatory cytokine and chemokine concentrations, viral load, and overall host morbidity, we sought to assess the severity of lung lesions as well (Fig. 4A-D). Histopathological analyses revealed extensive histopathologic changes in the lungs of OVA-sensitized mice compared to PBS controls, most prominent during the acute phase of inflammation (Fig. 4A-B, S6). Significant lesions were observed from 1 DPC through 7 DPC, before decreasing at 10 DPC. OVA-sensitized mice still had significantly higher total pathology scores compared to control mice at 28 DPC. Extensive neutrophil, eosinophil, lymphocyte, plasma cell, and macrophage inflammation were observed in OVA-sensitized mice, corroborating flow cytometry results (Fig. S6B-C). Mast cells and basophils were not substantially observed, likely due to cell rarity and the specific portion of the lung evaluated; mast cells are more prevalent in the upper respiratory tract rather than the lower respiratory tract, such as the lung lobes.

**Figure 4.**
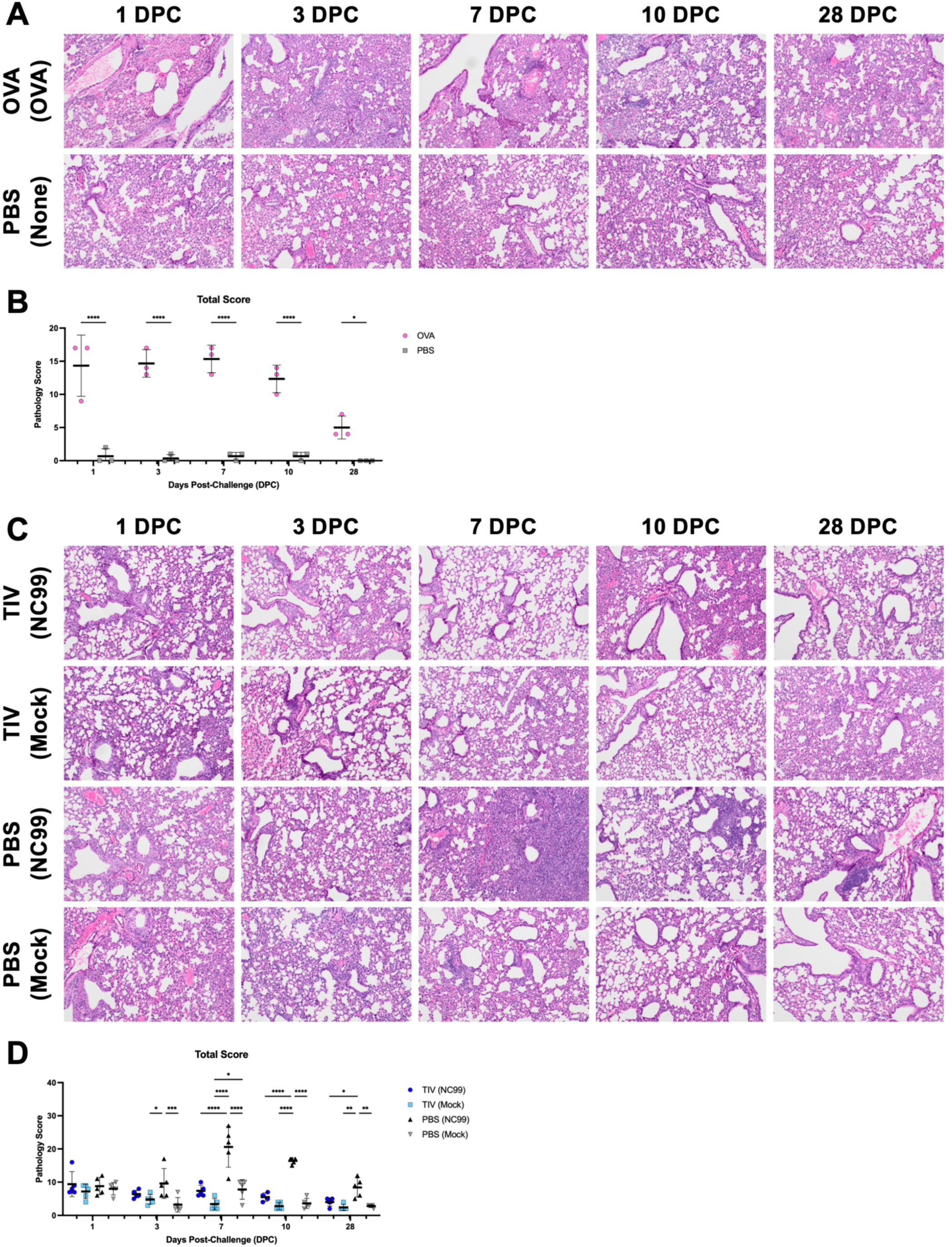
Breakthrough infection exhibits less disease compared to OVA allergic sensitization or primary influenza infection. Lung sections were stained with H&E then subjected to blinded pathology scoring. (**A**) Representative images and (**B**) total pathology score for the OVA sensitization model. (**C**) Representative images and (**D**) total pathology score for the breakthrough infection model. The total pathology score was comprised of the sum of the following metrics, individually scored from a scale of 1 to 5+: amount of lung affected and perivascular inflammation within the total lung; neutrophil inflammation, eosinophil inflammation, basophil inflammation, mast cell inflammation, lymphocyte/plasma cell inflammation, epithelial necrosis, intraluminal cells or debris, bronchus associated lymphoid tissue (BALT) hyperplasia in the bronchi and bronchioles or peribronchial and peribronchiolar areas; neutrophil inflammation, eosinophil inflammation, basophil inflammation, mast cell inflammation, lymphocyte/plasma cell/macrophage inflammation, necrosis/fibrin, consolidation, and edema in the alveoli or alveolar septa. Lines and error bars represent mean ± SD, with each symbol representing an individual mouse. Statistical significance was determined via ordinary two-way ANOVA with (**B**) Šídák’s multiple comparisons test with a single pooled variance, or (**D**) Tukey’s multiple comparisons test with a single pooled variance. ****P < 0.0001, ***P = 0.0001 to 0.001, **P = 0.001 to 0.01, *P = 0.01 to 0.05.

Severe lung lesions were observed in the PBS-vaccinated NC99-challenged primary infection mice when compared to other groups, such as the breakthrough infection mice and mock-challenged controls (Fig. 4C-D, S7). General patterns for neutrophil, eosinophil, and macrophage inflammation observed in histopathological analyses were in line with findings from flow cytometry (Fig. S7B-C). Scores for eosinophilic inflammation were lower for mice in the breakthrough infection model compared to the OVA-sensitization model. Primary infection mice had severe disease, unlike the breakthrough infection mice, with significantly higher total scores than all other groups at 7-28 DPC. Epithelial necrosis, as well as intraluminal cells and debris were most prevalent in primary infection mice compared to breakthrough infection mice and mock-challenged controls (Fig. S7B). Bronchus associated lymphoid tissue (BALT) hyperplasia was only observed in primary infection mice, beginning from 7 DPC and scores steadily increased through 28 DPC (Fig. S7B). Primary infection mice also had substantial consolidation, necrosis, and fibrin in the alveoli and alveolar septa at 7-10 DPC while other groups did not score highly for these metrics (Fig. S7C). Breakthrough infection mice that received vaccination and had a lower viral load did have some histopathologic changes, but not to the same extent as primary infection mice: higher perivascular inflammation and epithelial necrosis were observed in TIV-vaccinated NC99-challenged breakthrough infection mice relative to TIV-vaccinated mock-challenged controls, particularly at 1-10 DPC (Fig. S7A-B). However, pathology scores were significantly higher in the PBS-vaccinated NC99-challenged group compared to the TIV-vaccinated NC99-challenged group for the same metrics.

To better understand host health, we then employed another metric of lung pathology. Excess mucin has been associated with Type 2-skewed disease states, such as allergy, asthma, and RSV VAERD.^18^ As such, we scored Alcian Blue/Periodic acid–Schiff stained sections to determine the degree of mucin and goblet cell hyperplasia. We observed extensive mucin staining and goblet cell hyperplasia in OVA-sensitized, allergic mice most prevalently at 7 and 10 DPC (Fig. 5A-B). No mucin staining or goblet cell hyperplasia was observed in PBS controls. No substantial mucin staining or goblet cell hyperplasia was observed for any of the treatment groups in the breakthrough influenza infection model as well (Fig. 5C-D).

**Figure 5.**
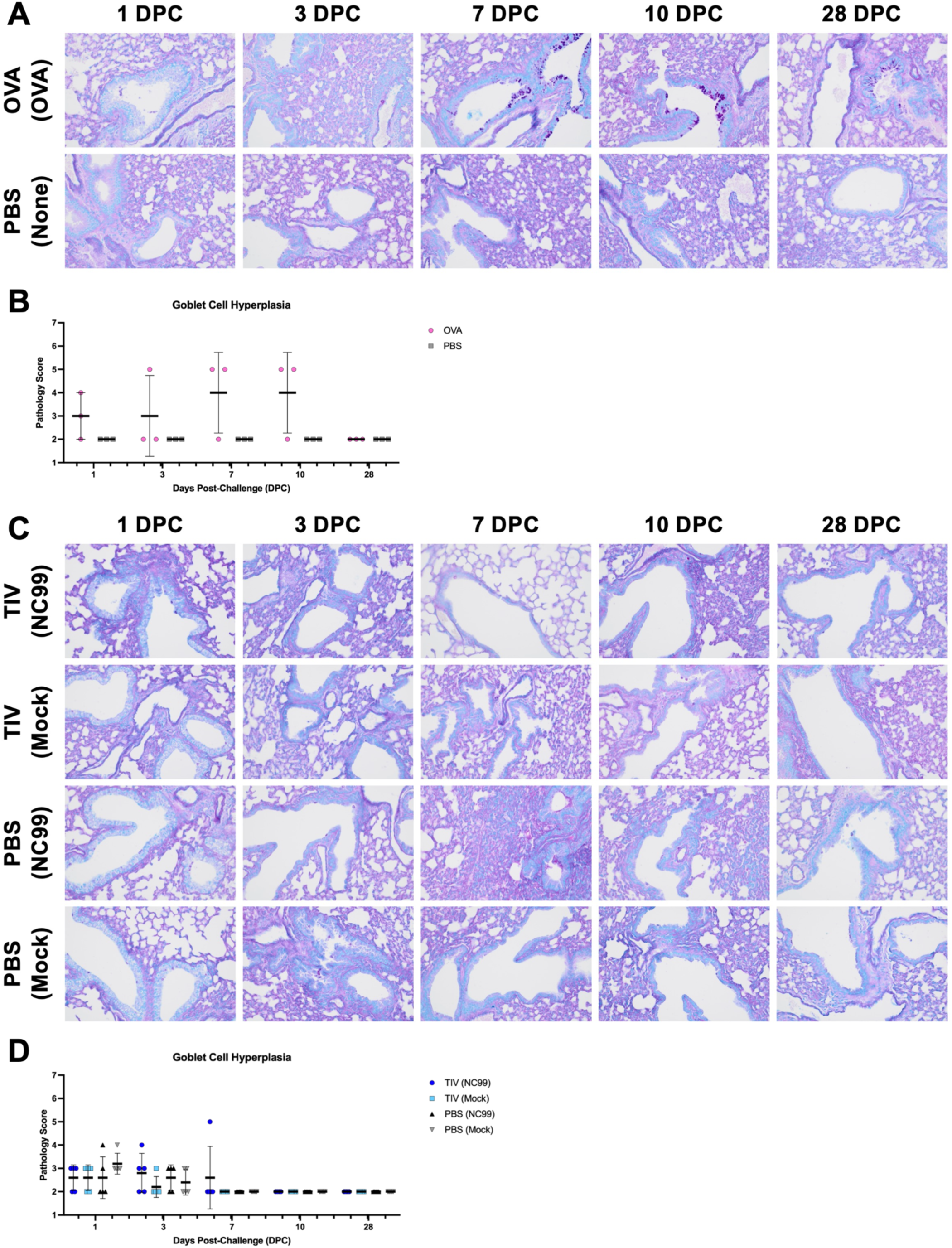
Goblet cell hyperplasia and positive mucin staining is observed during Type 2 allergic responses but not during breakthrough or primary influenza infection. Lung sections were stained with AB-PAS to assess mucin staining and goblet cell hyperplasia, then subjected to binded pathology scoring. (**A**) Representative images and (**B**) goblet cell score for the OVA sensitization model. (**C**) Representative images and (**D**) goblet cell score for the breakthrough infection model. Lines and error bars represent mean ± SD, with each symbol representing an individual mouse. Statistical significance was determined via ordinary two-way ANOVA with (**B**) Šídák’s multiple comparisons test with a single pooled variance, or (**D**) Tukey’s multiple comparisons test with a single pooled variance.

### OVA-sensitization and primary influenza infection result in isotype-skewed serum antibody responses, while TIV-vaccination confers a balanced IgG2a/IgG1 profile

We quantified total antigen-specific IgG, IgG1, and IgG2a titers to evaluate host humoral immunity after vaccination and challenge. In OVA-sensitized mice, we had detectable titers of OVA-specific total IgG by 2 weeks post-vaccination, which continued to steadily rise post-challenge (Fig. 6A). OVA-specific IgG titers were negligible for PBS control mice throughout the entire study. Similar patterns were observed for both treatment groups regarding OVA-specific IgG1 (Fig. 6B). Given the mice were primed with two intraperitoneal doses of OVA with alum adjuvant, a Type 2-skewing adjuvant^46^, we did not expect to see substantial titers of IgG2a in OVA-sensitized mice. Indeed, OVA-specific IgG2a titers were only detectable at 28-30 DPC with intranasal OVA (Fig. 6C). We used the ratio of antigen-specific IgG2a to IgG1 titers as a surrogate for evaluating Th1/Th2-skewing, with higher values indicating more IgG2a- and Th1-biased responses and lower values reflecting IgG1- and Th2-biased responses.^47–49^ OVA-sensitized mice exhibited very low IgG2a/IgG1 ratios, due to the heavily skewed IgG1 response and in line with the dogmatic Type 2 immune profile in the lung (Fig. 6D). Furthermore, OVA-sensitized mice also generated increasingly higher concentrations of total IgE following intranasal OVA challenge, with peak concentrations observed at 28-30 DPC, significantly greater than concentrations in PBS control mice (Fig. 6E).

**Figure 6.**
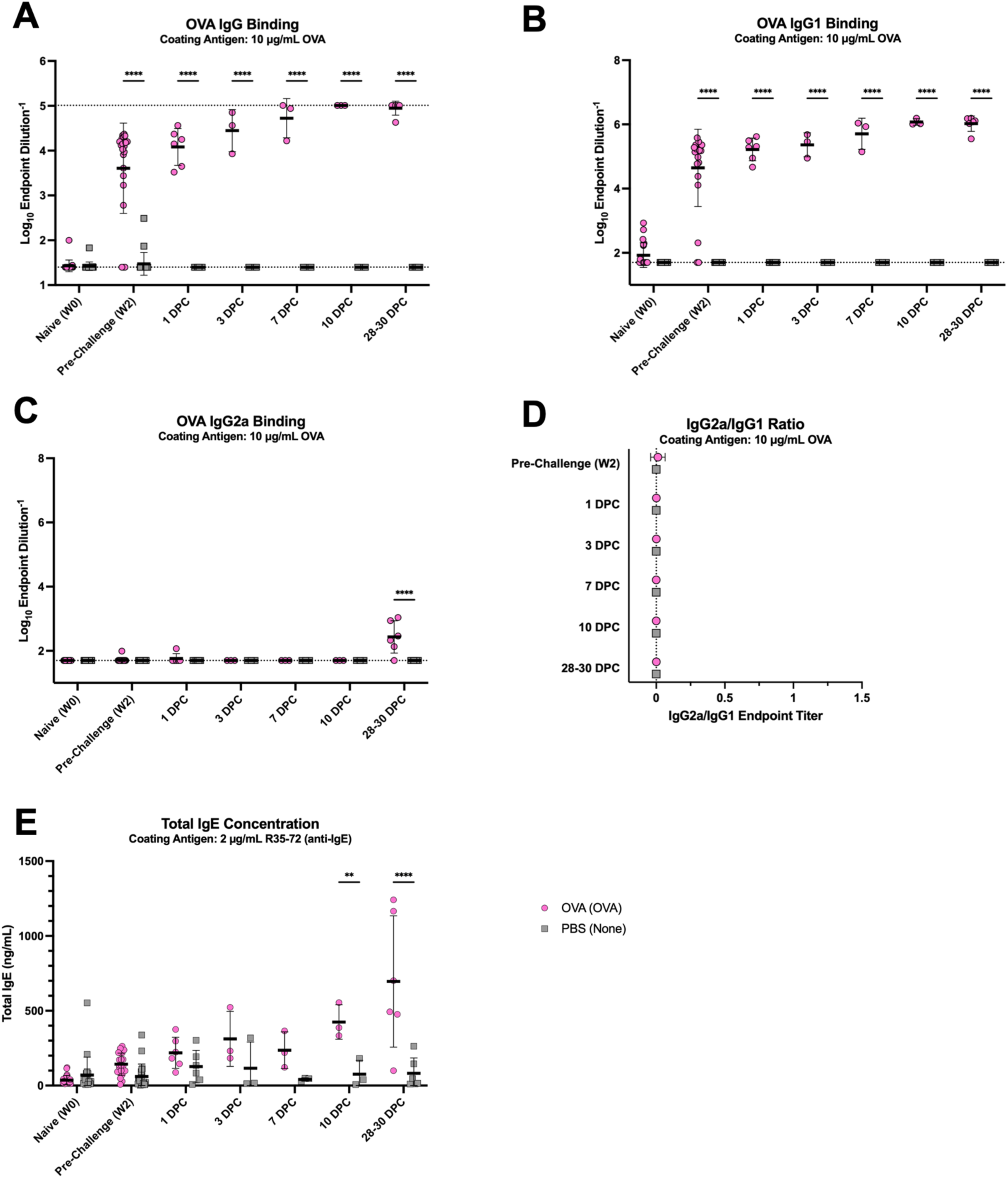
OVA-sensitized mice mount IgG1-biased serum antibody responses with increasing total IgE concentrations after intranasal challenge with antigen. OVA-binding (**A**) total IgG, (**B**) IgG1, and (**C**) IgG2a endpoint titers were measured. Dotted line represents an endpoint titer LOD of (**A**) 25 or (**B-C**) 50. (**A**) Total OVA IgG binding had an upper LOD of 102400. Undetectable values were arbitrarily set to the LOD. (**D**) The ratio of IgG2a/IgG1 endpoint titers was used to evaluate immune skewing. Dotted line represents a ratio of 0. Mice that had undetectable IgG1 and IgG2a endpoint titers for a given time point were arbitrarily set to a ratio of 0. (**E**) Total serum IgE concentrations were measured. Lines and error bars represent mean ± SD, with each symbol representing an individual mouse. Statistical significance was determined via ordinary two-way ANOVA with Šídák’s multiple comparisons test with a single pooled variance. ****P < 0.0001, **P = 0.001 to 0.01.

In the breakthrough infection study, we measured vaccine-specific total IgG, IgG1, and IgG2a titers similar to analyses done for the OVA sensitization mice. High titers of TIV-specific total IgG were detected in the two vaccinated groups at 2 weeks post-vaccination (Fig. 7A). TIV-specific total IgG titers continued to rise post-challenge for both TIV-vaccinated groups, although NC99-challenged mice had significantly higher titers than the mock-challenged group at 10 DPC, suggesting a systemic boosting effect from encountering viral antigen in the respiratory tract. TIV-specific IgG titers remained undetectable in PBS-vaccinated mice until 10 DPC following influenza infection, indicative of a *de novo* immune response against NC99. PBS-vaccinated NC99-challenged primary infection mice had higher TIV-specific titers by 28 DPC, but still had significantly lower titers compared to mice that were TIV-vaccinated. Similar patterns were observed for both IgG1 and IgG2a, although both isotypes were not as detectable for the primary infection mice at 10 DPC (Fig. 7B-C). Additionally, TIV-specific IgG2a titers were significantly higher in TIV-vaccinated NC99-challenged breakthrough infection mice compared to TIV-vaccinated mock-challenged controls, but this difference was not statistically significant for TIV-specific IgG1 titers. Given the similar increases in both IgG1 and IgG2a titers following vaccination and challenge, all TIV-vaccinated mice exhibited balanced IgG2a/IgG1 ratios through 3 DPC (Fig. 7D). At 7 and 10 DPC, the breakthrough infection mice had more skewing towards IgG2a relative to mock-challenged controls, although this skewing was transient and the ratios resembled the mock-challenged controls by 28 DPC. The PBS-vaccinated NC99-challenged mice undergoing a primary influenza infection, dogmatically Type 1-biased event, had a significantly IgG2a-skewed response that was evident at 28 DPC. There were transiently higher concentrations of total serum IgE in breakthrough infection mice at 3 DPC, although not as high as measurements in the OVA-sensitized, allergic mice (Fig. 7E).

**Figure 7.**
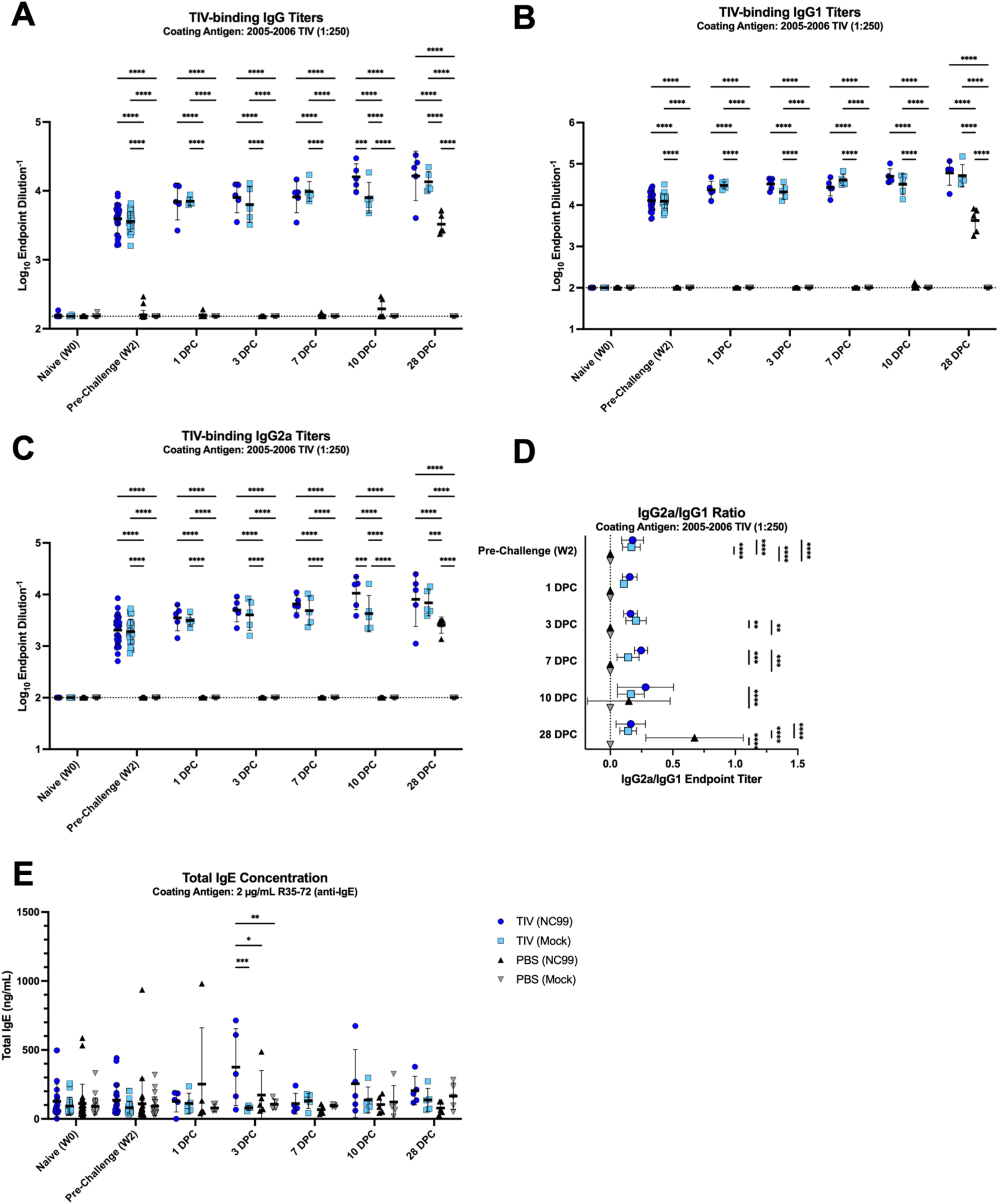
TIV-vaccinated mice seroconvert by 2 weeks post-vaccination and have balanced IgG1 and IgG2a titers with some increase in total IgE concentration shortly after sublethal viral challenge, while primary infection mice have IgG2a-biased responses following challenge. TIV-binding (**A**) total IgG, (**B**) IgG1, and (**C**) IgG2a endpoint titers were measured. Dotted line represents an endpoint titer LOD of (**A-C**) 100. Undetectable values were arbitrarily set to the LOD. (**D**) The ratio of IgG2a/IgG1 endpoint titers was used to evaluate immune skewing. Dotted line represents a ratio of 0. Mice that had undetectable IgG1 and IgG2a endpoint titers for a given time point were arbitrarily set to a ratio of 0. (**E**) Total serum IgE concentrations were measured. Lines and error bars represent mean ± SD, with each symbol representing an individual mouse. Statistical significance was determined via ordinary two-way ANOVA with Tukey’s multiple comparisons test with a single pooled variance. ****P < 0.0001, ***P = 0.0001 to 0.001, **P = 0.001 to 0.01, *P = 0.01 to 0.05.

Overall, our analyses of the antigen-specific response in the serum was in line with lung immune cell quantification, cytokine and chemokine measurement, and histopathology. The overt Type 2 bias observed in OVA-sensitized mice was corroborated by high OVA-specific IgG1 titers with little to no IgG2a. Similarly, the Type 1 biased lung response in primary influenza infection mice was mirrored in the IgG2a-skewed serum responses to antigen at later time points post-challenge. Breakthrough infection mice had balanced IgG2a/IgG1 ratios, indicative of a balanced Type 1/2 response.

### Principal component analysis of multiple host immune metrics shows distinct, balanced Type 1/2 response for breakthrough infection mice

Given the large number of individual-level data collected for each mouse in these studies, we used principal component analysis (PCA) to visualize differences between treatment groups and time points (Fig. 8). We integrated a total of 67 parameters: immune cell counts from flow cytometry, body weight percentages, cytokine and chemokine concentrations, histopathology scores, mucin staining scores, and total IgE concentrations. Antigen-specific antibody measurements or influenza-specific assay results were omitted to allow for head-to-head comparison of all treatment conditions at once. Distinct profiles for primary infection, breakthrough infection, and OVA-sensitization were observed. Differences in immune profile by time point could also be observed: the primary infection mice at 7 DPC and OVA-sensitized mice at 1 DPC varied the most along PC1 and PC2, respectively, compared to mock-challenged controls. Additionally, later time points such as 28 DPC for both conditions clustered more closely to mock-challenged controls. Primary infection mice varied the most on principal component 1 (PC1), driven by metrics such as body weight loss, AM number, and IL-33 concentration (Fig. S8). OVA-sensitized mice varied the most on principal component 2 (PC2), driven by eosinophil counts via flow cytometry and pathology, and concentrations of canonical Type 2 cytokines such as IL-4 and IL-5. Breakthrough infection mice had an intermediate immune profile between the primary infection and OVA-sensitized mice, clustering close to mock-challenged controls.

**Figure 8.**
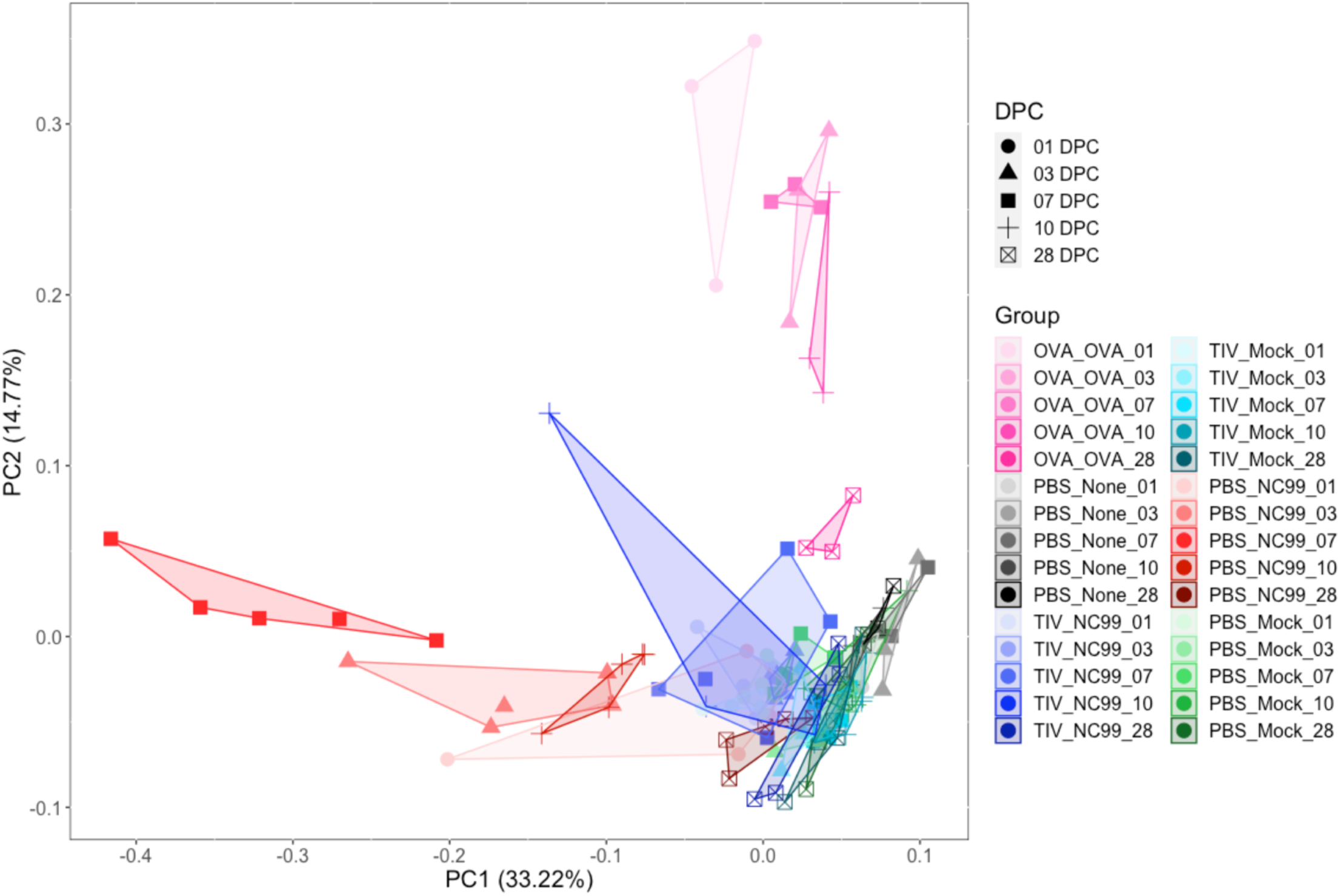
Principal component analysis confirms balanced systemic host immune response in breakthrough infection mice relative to primary infection and OVA mice. A total of 67 different immune metrics from each mouse was used for principal component analysis (PCA): the counts of 5 immune cell populations, body weight percentages from up to 12 time points, concentrations of 27 cytokines/chemokines, pathology scores for 19 different metrics, goblet cell scores, and total IgE concentration from 3 time points. Antigen-specific IgG, IgG1, IgG2a, IgG2a/IgG1 ratios, and TCID_50_ values were excluded from analysis to generate antigen-agnostic immune profiles. Symbol colors denote treatments, and symbol shapes denote time point post-challenge. Each symbol represents an individual mouse.

### Imaging confirms cellular dynamics seen by flow cytometry, identifies cell-cell interactions in lung space, and visualizes granulocyte extracellular trap formation

We used multiplex fluorescence imaging to visualize our immune cell populations of interest and understand cell-cell interactions for all models used (Fig. 9A). For this study, we included all four breakthrough infection model groups, as well as an OVA-sensitized group as a positive control for lung eosinophilia and a naive, negative control group as baseline. Based on data from our kinetics studies, we moved forward with 7 DPC as our time point of interest since our primary focus is in lung eosinophils during breakthrough infection, with a special emphasis on Siglec-F^hi^ eosinophils. Although total eosinophil numbers were the highest at 10 DPC, we selected 7 DPC due to more substantial enrichment for the Siglec-F^hi^ subset at this time point.

**Figure 9.**
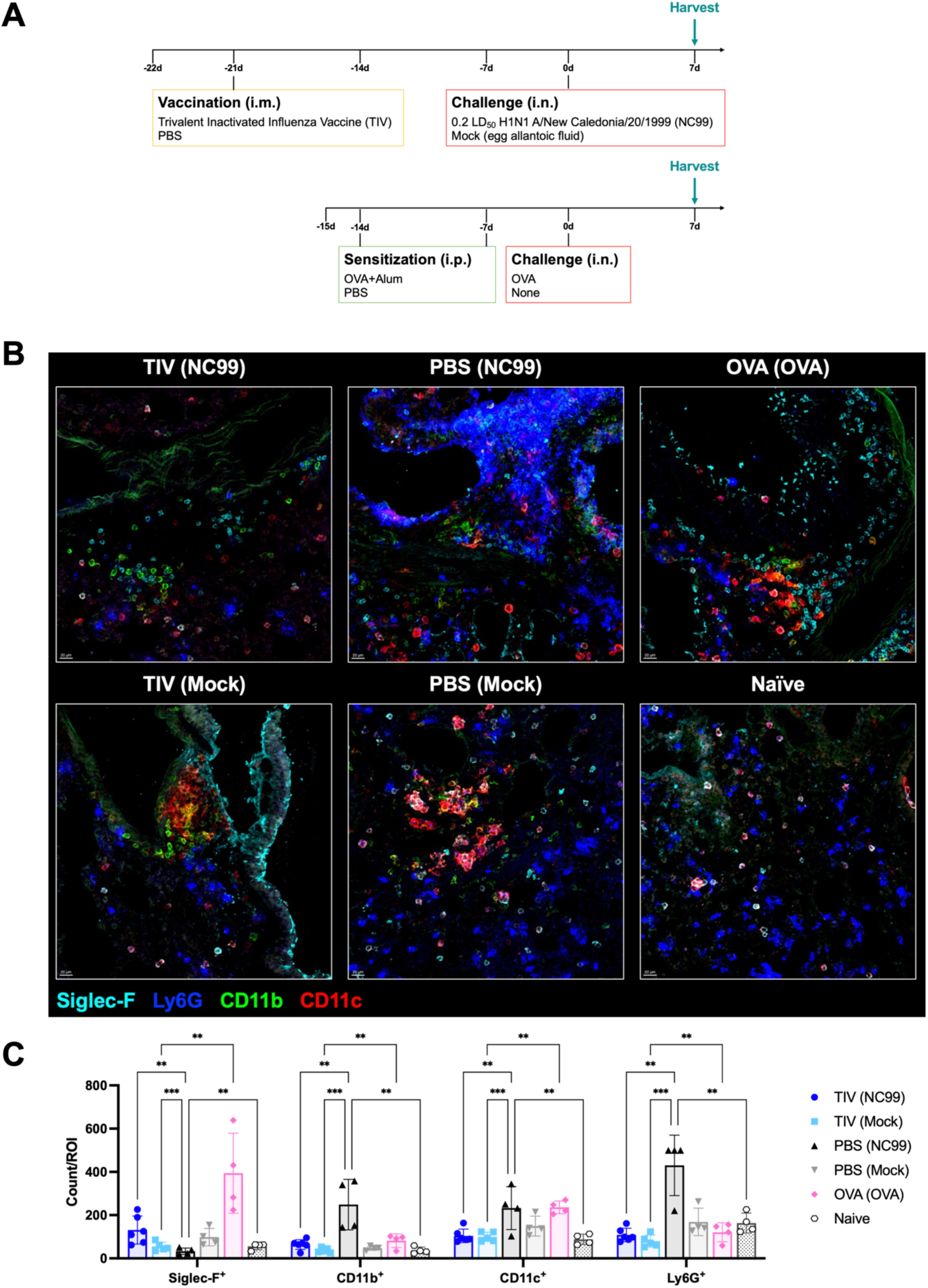
Multiparameter confocal microscopy confirms immune cell dynamics observed by flow cytometry in the lung at 7 DPC. Lung sections were stained for Siglec-F, CD11b, CD11c, and Ly6G to visualize major myeloid populations. (**A**) Representative images and (**B**) number of marker-positive cells was quantified for each region of interest (ROI). Bar plots and error bars represent mean ± SD, with each symbol representing an individual mouse. Scale bar in the bottom left corner of each image represents 20 µm. Statistical significance was determined via ordinary two-way ANOVA with main effects only, and with Tukey’s multiple comparisons test using a single pooled variance. ***P = 0.0001 to 0.001, **P = 0.001 to 0.01.

First, we quantified the number of Siglec-F^+^ (eosinophils and AMs), CD11b^+^ (macrophages), CD11c^+^ (dendritic cells), and Ly6G^+^ (neutrophils) cells using microscopy (Fig. 9B-C). In line with findings from flow cytometry, we observed a significantly higher number of Siglec-F^+^ cells per region of interest (ROI) in breakthrough infection mice and OVA-sensitized mice compared to all other groups (Fig. 9C). As seen in flow cytometry, the PBS-vaccinated NC99-challenged primary influenza infection mice had significantly higher counts of Ly6G^+^ cells compared to other groups. The primary influenza infection group also had higher counts per ROI of CD11b^+^ cells and CD11c^+^ cells.

Next, we stained lung sections for Siglec-F (eosinophils and AMs), Ly6G (neutrophils), CD11c (dendritic cells), CD3 (T cells), and B220 (B cells) to visualize granulocyte-lymphocyte interactions in the lung (Fig. 10A). We also included CD101 to identify inflammatory, activated eosinophils. Building upon data from the previous imaging panel and our flow cytometry data, we also observed a significantly higher number of CD3^+^ and CD101^+^ cells in the lungs of PBS-vaccinated NC99-challenged primary infection mice (Fig. 10B). We observed a multitude of interactions between Siglec-F^+^ cells and Ly6G^+^ cells with CD3^+^ cells (Fig. 10A). Of note, we noticed multiple areas of CD101^+^ Siglec-F^+^ cells interacting with CD3^+^ cells in the breakthrough infection and OVA-sensitized lungs, potentially indicating activated eosinophils interfacing with T cells (Fig. S9).

**Figure 10.**
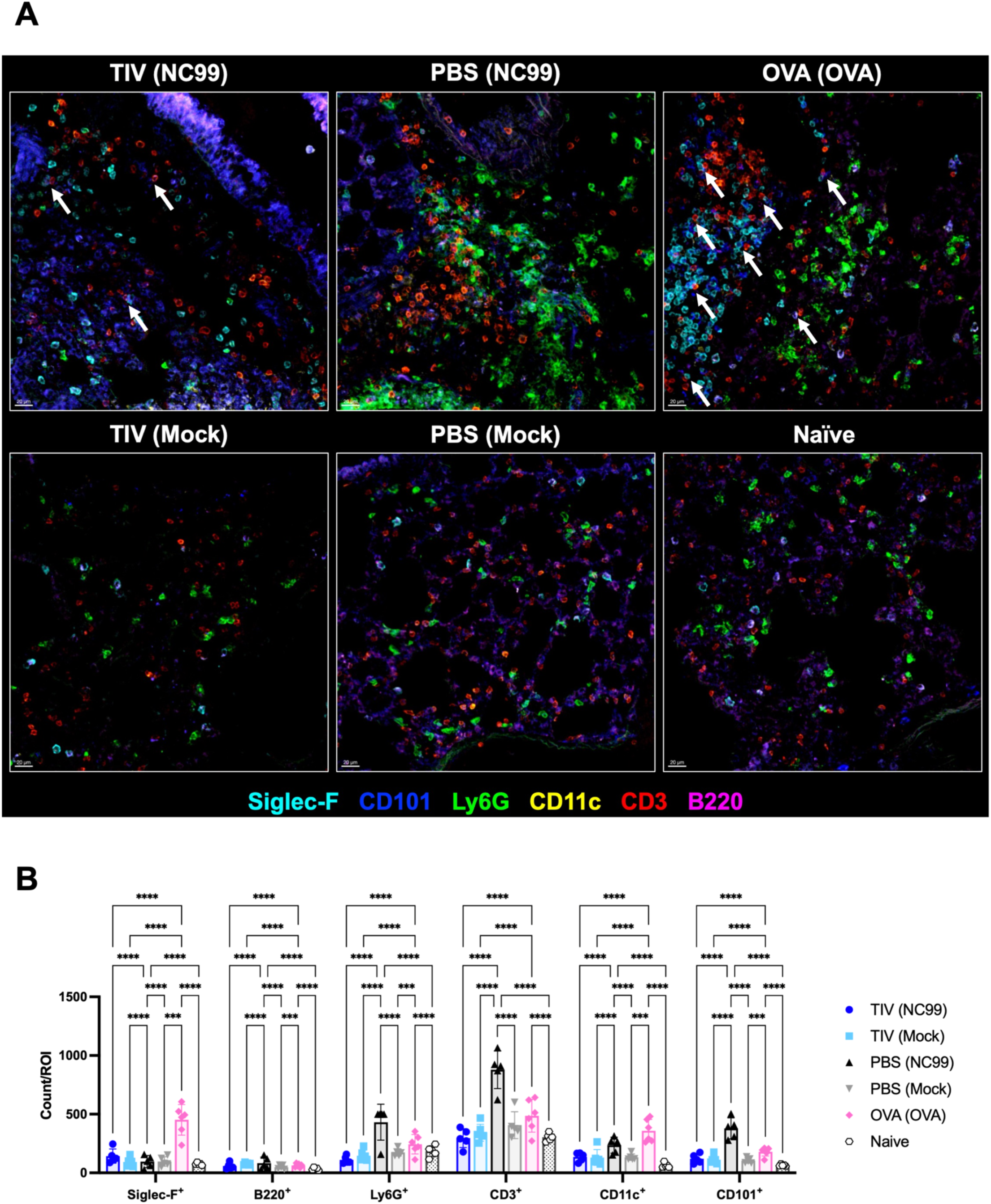
Granulocyte-T interactions can be visualized in the lung parenchymal space following breakthrough infection, primary infection, or allergic sensitization. Lung sections were stained for Siglec-F, CD101, Ly6G, CD11c, CD3, and B220 to quantify cell-cell interactions in tissue. (**A**) Representative images and (**B**) number of marker-positive cells was quantified for each region of interest (ROI). White arrows denote regions of CD101^+^ Siglec-F^+^ cells interacting with CD3^+^ cells. Bar plots and error bars represent mean ± SD, with each symbol representing an individual mouse. Scale bar in the bottom left corner of each image represents 20 µm. Statistical significance was determined via ordinary two-way ANOVA with main effects only, and with Tukey’s multiple comparisons test using a single pooled variance. ****P < 0.0001, ***P = 0.0001 to 0.001.

Since neutrophil or eosinophil extracellular traps (NETs and EETs, respectively) can exacerbate host lung disease during respiratory viral infection or asthma, we stained lung sections to check for the presence of NETs or EETs (Fig. 11). NETs have some properties that aid in clearing viral infection, such as direct antiviral activity against influenza, conferred by arginine-rich histones H3 and H4 released during NET formation.^50,51^ However, NETs have also been shown to damage lung tissue, promote inflammation, and decrease gas exchange, thereby significantly increasing disease severity.^51–53^ EETs have been predominantly described in the context of extracellular pathogens, such as bacteria, and implicated in the pathogenesis of asthma.^54–56^ We also stained sections for Siglec-F^+^ cells and Ly6G^+^ cells, as well as a DNA stain to identify nuclei (DRAQ7^+^). Qualitatively, we observed Siglec-F^+^ cells in lungs of TIV-vaccinated NC99-challenged breakthrough infection mice, but in the absence of cell-free eosinophil peroxidase (EPX). This is in stark contrast with the OVA-sensitized mice, which had multiple areas with extensive cell-free EPX and citrullinated H3 staining, indicative of EET and NET formation, respectively.^57,58^ NET formation was also prominently observed in PBS-vaccinated NC99-challenged primary influenza infection mice. NET formation in the lungs following viral challenge has been described for primary SARS-CoV-2 infection.^59^

**Figure 11.**
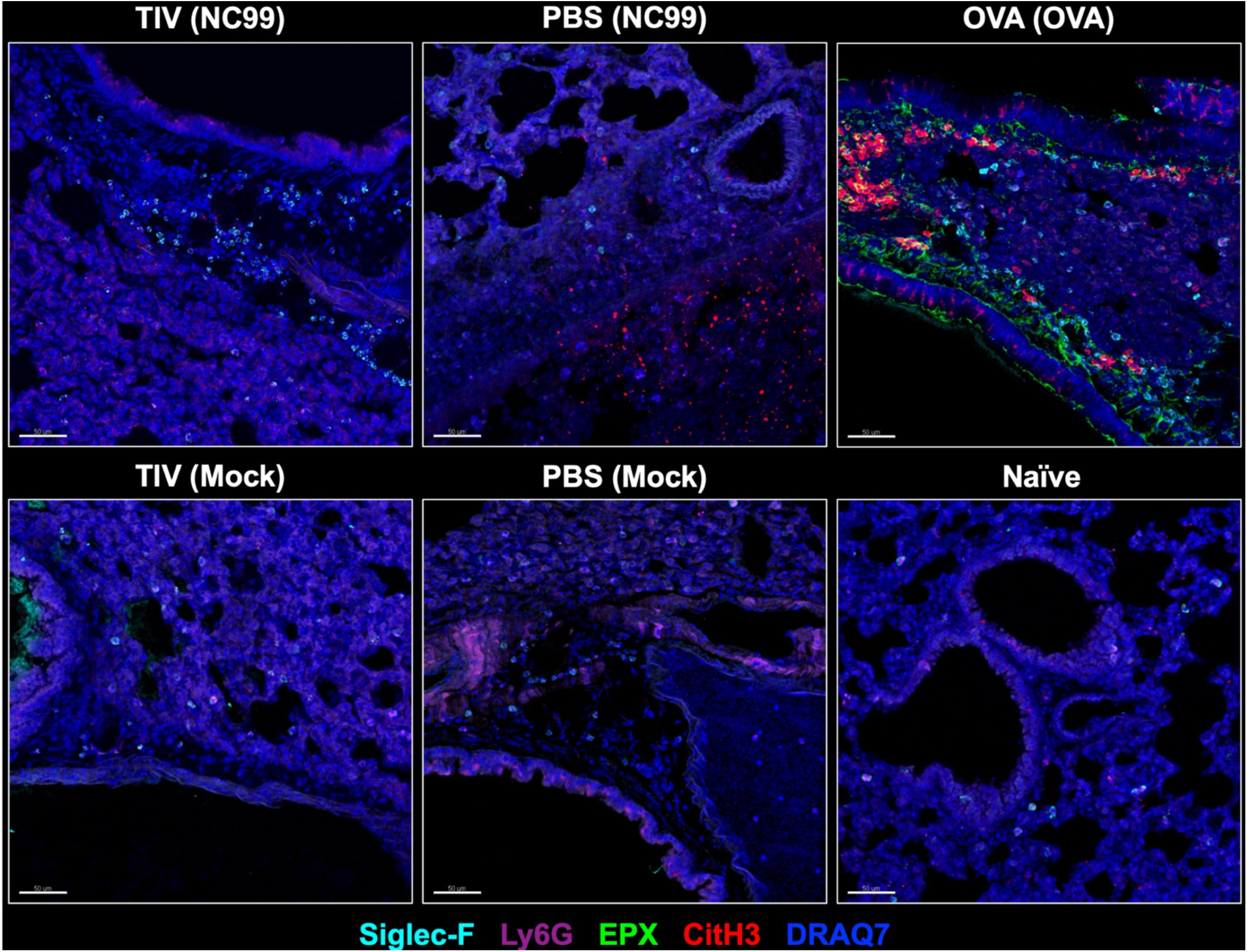
Extracellular trap release is most prominently seen after allergic sensitization or primary infection, but not breakthrough infection. Lung sections were stained with Siglec-F, Ly6G, EPX, and citrullinated H3 antibodies alongside a DNA stain, DRAQ7, to evaluate neutrophil (CitH3^+^) and eosinophil extracellular trap (cell-free EPX^+^) formation in tissue. Scale bar in the bottom left corner of each image represents 50 µm. Representative images of each treatment group are shown here.

## Discussion

This study provides a comprehensive, longitudinal analysis of immune responses in the lungs across three distinct conditions: OVA sensitization as a typical Type 2 response, primary influenza infection as a Type 1 response, and breakthrough influenza infection in vaccinated hosts as a mixed Type 1/2 response. By examining the kinetics of immune cell populations, cytokine and chemokine profiles, and lung pathology, critical differences in the dynamics and consequences of these responses were revealed. Using imaging to evaluate differences in treatment groups at 7 DPC, a time point exhibiting high eosinophils in OVA-sensitized and breakthrough infection mice, we visualized granulocyte-T cell interactions in the lung tissue space alongside NET and EET formation.

OVA sensitization induced a classic Type 2 immune response characterized by robust eosinophilic infiltration, transient neutrophilia, and AM depletion during the acute phase. Eosinophil recruitment in OVA-sensitized mice was particularly dramatic, with a marked enrichment of the Siglec-F^hi^ subset, which is thought to be an activated, pro-inflammatory subset.^25,43–45^ This response was driven by elevated Type 2 cytokines, including IL-4, IL-5, and IL-13, which are known to promote goblet cell hyperplasia and mucin production, hallmark features of allergic inflammation.^41,60^ Findings in the lungs were corroborated by analysis of the antigen-specific humoral response: OVA-sensitized mice generated an IgG1-dominated OVA-specific response, which is indicative of Th2-skewing, and high concentrations of total IgE in the serum.^47–49^

In contrast, primary influenza infection elicited a strong Type 1 response dominated by neutrophilic inflammation and prolonged AM depletion. The cytokine profile was consistent with antiviral immunity, with high levels of IFN-γ, TNF-α, and IL-12. Unlike the OVA sensitization model, there was no substantial eosinophilia post-challenge, and significant lung pathology was observed. Viral load measurements indicated prolonged viral persistence in primary infection mice compared to vaccinated breakthrough infection mice, likely due to the lack of pre-existing immunity. Characterization of the humoral response post-challenge confirmed elicitation of a Th1-skewed response, with IgG2a being the dominant isotype.^47–49^

Breakthrough influenza infection in vaccinated mice presented a mixed Type 1/2 response, without the heavy Type 1 skewing observed in primary infection mice. Eosinophil recruitment was moderate but significant, with an enrichment of the Siglec-F^hi^ subset particularly at 7-10 DPC, although not to the extent observed in OVA-sensitized mice. Importantly, AM counts were preserved, and neutrophilic inflammation was mild, suggesting that vaccination mitigates excessive immune activation. Cytokine and chemokine levels in breakthrough infection mice were balanced, with muted Type 1 and Type 2 modules compared to primary infection and OVA sensitization, respectively. Surprisingly, concentrations of canonical Type 2 cytokines IL-4, IL-5, and IL-13 were highest in the primary infection mice, despite the lack of eosinophilia (Fig. S5). Overall, the balanced immune response in breakthrough infection mice coincided with rapid viral clearance and minimal lung disease, emphasizing the protective effects of pre-existing immunity conferred by vaccination.

Vaccination significantly altered the immune landscape in breakthrough infection, preserving AM populations and preventing the severe neutrophilic inflammation observed in primary influenza infection. AMs are essential for maintaining lung homeostasis and orchestrating immune responses, and their preservation likely contributed to the regulated inflammatory environment in vaccinated hosts. Moreover, it is to be expected that preservation of AMs could have allowed for faster recovery and clearance of damaged cells, a metric for a more rapid return to homeostasis. Additionally, the absence of NETs and EETs in breakthrough infection mice underscores the protective nature of vaccine-induced immunity. Extracellular traps are associated with tissue damage and can exacerbate disease in severe viral infections and allergic diseases, to the detriment of the host.^54–56^ Their absence in the vaccinated hosts highlights the efficacy of vaccination in preventing excessive inflammation. The rapid viral clearance observed in vaccinated mice underscores the benefits of pre-existing immunity. Reduced viral titers correlated with minimal weight loss and quicker recovery, emphasizing the ability of vaccination to mitigate disease severity. These findings align with clinical observations of reduced morbidity in vaccinated individuals experiencing breakthrough infections, providing strong evidence for the effectiveness of influenza vaccines.

These findings reinforce the distinction between eosinophilic recruitment in mouse models of breakthrough influenza infection and vaccine-associated enhanced respiratory disease (VAERD). VAERD is characterized by severe disease, including goblet cell hyperplasia and excessive Type 2 cytokine responses, none of which were observed in breakthrough infection. While eosinophilia was present in vaccinated mice, it was non-immunopathological and associated with reduced viral titers and minimal lung damage. This suggests that rather than exacerbating disease, eosinophils may contribute to host defense or tissue repair when excessive inflammation is not present in the local lung environment. The enrichment of Siglec-F^hi^ eosinophils, often implicated in allergic conditions, did not reach the pathological levels observed in OVA-sensitized mice. In line with the Local Immunity And/or Remodeling/Repair (LIAR) hypothesis^14^, we speculate that the Siglec-F^hi^ eosinophils observed during breakthrough infection may be functionally distinct from the Siglec-F^hi^ eosinophils observed during allergic inflammation, due to differences in the host lung microenvironment; OVA-sensitized mice were extremely polarized towards Type 2 responses, whereas breakthrough infection mice have a more balanced Type 1/2 response (Fig. 8). Whether or not the wealth of Type 2 cytokines, chemokines, and other factors confer substantial functional differences to eosinophils in the lungs of allergen-challenged mice versus breakthrough infection mice remains to be seen, particularly for the Siglec-F^hi^ subset. In depth phenotyping or transcriptomic analysis of eosinophils across multiple different inflammatory states will shed light on the differences between eosinophil subsets.

The specific role of eosinophils in breakthrough respiratory infection remains an area of interest. Traditionally associated with allergic inflammation, eosinophils have emerging roles in antiviral defense, such as antigen presentation and cytokine production.^9–13^ In this study, eosinophilia in vaccinated mice was correlated with rapid viral clearance and reduced inflammation, suggesting a protective function. We also observed Siglec-F^+^ cells interacting with CD3^+^ T cells, most prevalently in OVA-sensitized mice (Fig. 10, S9). Wiese et al. have also observed inflammatory eosinophils (Siglec-F^+^CCR3^+^CD11c^+^) interacting with T cells in the lungs of mice with allergic asthma, which were able to promote antigen-specific proliferation and differentiation of naive T cells.^61^ We also visualized activated, inflammatory eosinophils (CD101^+^ Siglec-F^+^ cells) in contact with T cells in the lung. It is possible that eosinophils were presenting antigen to T cells, although it is unlikely there were high levels of viral antigen present at the time point selected for imaging; by 7 DPC, viral titers are entirely controlled in breakthrough infection mice. Other possible eosinophil-T cell interactions of interest include immunomodulation.^12,62^ CD101 is associated with highly immunosuppressive Tregs and myeloid cells.^12,63–65^ Whether or not CD101^+^ eosinophils are able to suppress T cells has not been reported, but is a possible function. Other putative roles for eosinophils during breakthrough infection of respiratory viruses in vaccinated hosts have been reviewed by Chang and Schotsaert.^12^ However, the definitive roles of eosinophils during breakthrough infection or why they are recruited have yet to be defined. Future research should investigate the specific mechanisms by which eosinophils contribute to immune defense in this context and delineate the thresholds at which eosinophilia transitions from protective to pathological. We are currently investigating the precise contribution of eosinophils during breakthrough infection by leveraging eosinophil-specific depletion models or eosinophil-depleting antibodies, in conjunction with functional studies and single cell profiling.

One of the goals of our longitudinal study was to identify a recruitment signal for eosinophils during breakthrough infection via cytokine and chemokine profiling. We did not find a clear, conclusive recruitment signal for eosinophils in the breakthrough infection model. In general, most cytokine or chemokine expression patterns we observed in breakthrough infection mice were similar to that of primary infection mice, except substantially lower in concentration. We suspect this is because the strength of the cytokine/chemokine signal is directly linked to the viral titer; primary infection mice had significantly higher viral load while breakthrough infection mice only had detectable viral load through 3 DPC before rapidly controlling viral replication (Fig. 3C). Of the 27 cytokines and chemokines we evaluated, 3 analytes deviated from the viral titer-associated rise-fall kinetics: CCL11, IL-27, and IL-33. CCL11, also known as Eotaxin-1, is a potent eosinophil chemoattractant.^66^ Interestingly, CCL11 measurements proved inconclusive in the OVA-sensitization model, as OVA-sensitized mice did not have significantly higher concentrations than PBS control mice (Fig. S4). Similarly, CCL11 concentrations were highest in the breakthrough infection mice only at 1 DPC, much earlier than peak lung eosinophilia at 7-10 DPC (Fig. S5). At other time points, CCL11 concentrations in breakthrough mice was not significantly higher than any of the other groups. We expected to see classic rise-fall kinetics in the concentrations of CCL11 and IL-5 for the breakthrough infection mice, corresponding with the gradual influx of lung eosinophil counts, but kinetics for these two analytes did not appear clearly linked to eosinophil numbers. It is possible that the chemokines or cytokines were already bound by cells and not detectable as a measurable, free-floating protein in the lung homogenate supernatants. Next, IL-27 also had unique expression patterns across the different treatment groups we evaluated. IL-27 is an immunomodulatory cytokine produced by antigen-presenting cells in response to toll-like receptor (TLR) stimulation that acts on both innate and adaptive cells.^67^ It has been linked to attenuation of allergic airway diseases, such as by repressing group 2 innate lymphoid cells (ILC2) expansion, and has multiple antiviral roles as well.^68,69^ During influenza infection, IL-27 can promote transcription of interferon-stimulated genes like *Mx1*, increase the number of influenza-specific IFN-γ+ CD8 T cells, and ameliorate immunopathology by stimulating T cell production of IL-10.^69^ IL-27 concentrations were the most elevated in OVA-sensitized mice at 1 DPC and in primary infection mice at 7 DPC, time points that corresponded to peak inflammation and pathology (Fig. 4, S5). It is likely that IL-27 expression increased at these specific time points as a negative feedback mechanism to dampen inflammation. Of note, IL-27 concentrations were the lowest in the breakthrough infection mice at 3 DPC compared to all other groups, including mock-challenged controls, despite detectable viral titers at this time point (Fig. 3C, S5). We speculate that below a certain inflammatory threshold, IL-27 expression is actively suppressed: because the sublethal infection was well-controlled by factors such as pre-existing vaccine-elicited antibodies in the breakthrough infection mice, the magnitude of inflammation triggered was lower, resulting in less immunopathology and therefore less need to induce a marked anti-inflammatory host state. In line with this idea, BALT hyperplasia was more prevalent in primary infection mice, but not breakthrough infection mice; iBALT formation is correlated with tissue damage (Fig. S7B).^70^ Altogether, the absence of overt tissue damage in vaccinated hosts experiencing breakthrough infection, relative to unvaccinated hosts, resulted in muted pro- and anti-inflammatory signatures. Lastly, IL-33 was uniquely reduced in the primary infection mice (Fig. S5). IL-33 is an alarmin cytokine released in response to tissue damage that can potently activate Type 2 immune effector cells such as ILC2s.^71^ Our findings contrast with a previous report that primary influenza infection of C57BL/6 mice increases the number of IL-33 mRNA transcripts in the lungs; we observed a significant reduction of IL-33 protein, potentially highlighting mouse strain-specific differences in lung immunity since we worked with BALB/c mice.^72^ Of note, the kinetics of IL-33 concentrations in primary infected mice paralleled that of AM counts, gradually decreasing through 7-10 DPC to significantly lower levels compared to other groups. No such reduction in IL-33 or AMs was observed in breakthrough infection mice, which had vaccine protection from sustained viral replication. Whether or not IL-33 and AM survival are linked has yet to be described, although IL-33 has been shown to activate AMs in a mouse model of asthma.^73^ Collectively, although we did not identify a clear cytokine or chemokine recruitment signal for eosinophils into the lungs that neatly coincided with the timing of peak eosinophilia, it is possible that other factors like complement or lipid mediators are responsible for recruiting eosinophils in this scenario. Eosinophils can also be recruited via the C5a/C5aR1 axis^61^, serotonin metabolite 5-hydroxyindoleacetic acid (5-HIAA) binding to cell surface GPR35^74^, and arachidonic acid metabolites.^75^ Eotaxin-2 can also recruit eosinophils and was not within the cytokine/chemokine detection panel we used for these experiments, but will be included in future studies.^75^

Our study is believed to be the first to longitudinally profile breakthrough influenza infection across multiple time points and directly compare it to primary influenza infection and allergic sensitization in the lungs. Our detailed kinetic comparisons integrated flow cytometry, cytokine and chemokine analysis, morbidity, serology, imaging, and histopathology, providing a comprehensive, holistic view of immune dynamics. The high number of different immune metrics collected on an individual mouse basis allowed us to identify distinct immune profiles using unbiased methods. We also visualized novel CD101^+^Siglec-F^+^ cell and CD3^+^ T cell interactions, which we aim to investigate further in future studies. Additionally, we demonstrate that imaging of granulocyte extracellular traps can provide insight on host immunity and lung health: staining for both EET and NET formation in lung sections can be used to visually determine pathological granulocyte function.

Although our study has multiple strengths, certain limitations of the study warrant consideration. The lack of perfusion or intravascular staining before lung harvest may have influenced immune cell counts, as some of the numbers could reflect circulating cells rather than true tissue infiltration. However, CD101^+^ Siglec-F^hi^ eosinophils have been demonstrated to be in the lung tissue by multiple independent groups, as this population is predominantly protected from intravascular staining.^43,61,76,77^ Furthermore, we confirmed the enhanced presence of Siglec-F^+^ cells in the lung tissue via microscopy. Therefore we are confident that our quantification of Siglec-F^hi^ eosinophils reflects a lung-specific population, rather than a circulating population. Given the scope of the study, we did not include a post-prime pre-challenge lung harvest. The absence of a pre-challenge time point limits the ability to assess whether vaccination itself remodeled the lung immune environment. The effects of vaccination on tissues distal from the injection site, such as training and alterations of bone marrow progenitor cell populations, will be evaluated in future studies. Ultimately, we present a mouse model for breakthrough influenza infection in vaccinated hosts; confirmation that eosinophils are recruited to the lungs of humans experiencing breakthrough infection is warranted, perhaps through collection of bronchoalveolar lavage fluid or sputum. Continued surveillance of patients on eosinophil-depleting treatments and the effect of these treatments on their vaccine immune responses, particularly the incidence rate and severity of breakthrough infections in eosinophil-depleted patients, would be of great interest as well.

In conclusion, this study demonstrates that breakthrough influenza infection in vaccinated hosts elicits a balanced Type 1/2 immune response characterized by moderate eosinophilia, preserved AM counts, and minimal pathology. Unlike RSV VAERD, eosinophilia in breakthrough infection does not appear to be a cause for concern or a side effect of off-target vaccine immunity; this instance of lung eosinophilia was non-pathological and corresponded to enhanced immune defense. These findings underscore the protective role of vaccination in modulating immune responses, preventing excessive inflammation, and promoting rapid recovery. Our findings also highlight the potential for eosinophils to contribute positively to antiviral immunity under specific, balanced conditions. Our work has significant implications for understanding vaccine-induced immunity and optimizing vaccine strategies to achieve effective and safe protection against respiratory viral infections without enhancing immune pathology by clarifying that the induction of eosinophils is not always deleterious, and may not always warrant alarm.

## Data availability statement

The raw data supporting the conclusions of this article will be made available by the authors, without undue reservation.

## Ethics statement

The animal study was reviewed and approved by IACUC of the Icahn School of Medicine at Mount Sinai.

## Author contributions

LAC and MS conceptualized the project and research strategy. LAC, STY, BTW, PW, MN, and GS performed experiments, analyzed, and interpreted data. LAC and MS wrote the manuscript. All authors reviewed the manuscript and approved the submitted version.

## Funding

This research was supported in part by a Public Health Service Institutional Research T32 Training Award (AI07647) to LC. Influenza research in the MS lab is further supported by NIH/NIAID R21AI151229, R21AI176069, R44AI176894, CRIPT (Center for Research on Influenza Pathogenesis and Transmission), an NIH NIAID funded Center of Excellence for Influenza Research and Response (CEIRR, contract number 75N93021C00014) and by the NIAID funded SEM-CIVIC consortium to improve influenza vaccines (contract number 75N93019C00051).

## Conflict of interest

The authors declare this research was conducted in the absence of any commercial or financial relationships that could be construed as a potential conflict of interest. The MS laboratory has received unrelated research funding from sponsored research agreements from ArgenX BV, Moderna, 7Hills Pharma, and Phio Pharmaceuticals.

## Acknowledgements

We thank Eleanor Burgess and Johanna Vandekerckhove for feedback on initial drafts of the manuscript; Farah El-Ayache and Leonie Gruneberg for laboratory management; Daniel Flores, Ryan Camping, and Jane Deng for administrative support; Richard Cadagan for technical support; and the Mount Sinai Center for Comparative Medicine and Surgery for mouse colony management. We would also like to thank Dr. Elizabeth Jacobsen for providing the anti-EPX antibody.

The data in this paper were used in a dissertation as partial fulfillment of the requirements for a PhD degree at the Graduate School of Biomedical Sciences at Mount Sinai.

## References

1. Lambert LC, Fauci AS. Influenza vaccines for the future. N Engl J Med. 2010;363:2036–2044.

2. Cox NJ, Subbarao K. Influenza. Lancet. 1999;354:1277–1282.

3. Nichol KL, Treanor JJ. Vaccines for seasonal and pandemic influenza. J Infect Dis. 2006;194 Suppl 2:S111–8.

4. Osterholm MT, Kelley NS, Sommer A, et al. Efficacy and effectiveness of influenza vaccines: a systematic review and meta-analysis. Lancet Infect Dis. 2012;12:36–44.

5. Ferdinands JM, Thompson MG, Blanton L, et al. Does influenza vaccination attenuate the severity of breakthrough infections? A narrative review and recommendations for further research. Vaccine. 2021;39:3678–3695.

6. Grant EJ, Quiñones-Parra SM, Clemens EB, et al. Human influenza viruses and CD8(+) T cell responses. Curr Opin Virol. 2016;16:132–142.

7. Iwasaki A, Pillai PS. Innate immunity to influenza virus infection. Nat Rev Immunol. 2014;14:315–328.

8. Hashimoto Y, Moki T, Takizawa T, et al. Evidence for phagocytosis of influenza virus-infected, apoptotic cells by neutrophils and macrophages in mice. J Immunol. 2007;178:2448–2457.

9. Rosenberg HF, Dyer KD, Domachowske JB. Respiratory viruses and eosinophils: exploring the connections. Antiviral Res. 2009;83:1–9.

10. LeMessurier KS, Samarasinghe AE. Eosinophils: Nemeses of pulmonary pathogens? Curr Allergy Asthma Rep. 2019;19:36.

11. Gazzinelli-Guimaraes PH, Jones SM, Voehringer D, et al. Eosinophils as modulators of host defense during parasitic, fungal, bacterial, and viral infections. J Leukoc Biol. 2024;116:1301–1323.

12. Chang LA, Schotsaert M. Ally, adversary, or arbitrator? The context-dependent role of eosinophils in vaccination for respiratory viruses and subsequent breakthrough infections. J Leukoc Biol. 2024;116:224–243.

13. Samarasinghe AE, Melo RCN, Duan S, et al. Eosinophils Promote Antiviral Immunity in Mice Infected with Influenza A Virus. J Immunol. 2017;198:3214–3226.

14. Lee JJ, Jacobsen EA, McGarry MP, et al. Eosinophils in health and disease: the LIAR hypothesis. Clin Exp Allergy. 2010;40:563–575.

15. Chusid MJ. Eosinophils: Friends or foes? J Allergy Clin Immunol Pract. 2018;6:1439–1444.

16. Kim HW, Canchola JG, Brandt CD, et al. Respiratory syncytial virus disease in infants despite prior administration of antigenic inactivated vaccine. Am J Epidemiol. 1969;89:422–434.

17. Munoz FM, Cramer JP, Dekker CL, et al. Vaccine-associated enhanced disease: Case definition and guidelines for data collection, analysis, and presentation of immunization safety data. Vaccine. 2021;39:3053–3066.

18. Knudson CJ, Hartwig SM, Meyerholz DK, et al. RSV vaccine-enhanced disease is orchestrated by the combined actions of distinct CD4 T cell subsets. PLoS Pathog. 2015;11:e1004757.

19. Castilow EM, Legge KL, Varga SM. Cutting edge: Eosinophils do not contribute to respiratory syncytial virus vaccine-enhanced disease. J Immunol. 2008;181:6692–6696.

20. Rajão DS, Chen H, Perez DR, et al. Vaccine-associated enhanced respiratory disease is influenced by haemagglutinin and neuraminidase in whole inactivated influenza virus vaccines. J Gen Virol. 2016;97:1489–1499.

21. Kimble JB, Wymore Brand M, Kaplan BS, et al. Vaccine-Associated Enhanced Respiratory Disease following Influenza Virus Infection in Ferrets Recapitulates the Model in Pigs. J Virol. 2022;96:e0172521.

22. Khurana S, Loving CL, Manischewitz J, et al. Vaccine-induced anti-HA2 antibodies promote virus fusion and enhance influenza virus respiratory disease. Sci Transl Med. 2013;5:200ra114.

23. Choi A, Ibañez LI, Strohmeier S, et al. Non-sterilizing, Infection-Permissive Vaccination With Inactivated Influenza Virus Vaccine Reshapes Subsequent Virus Infection-Induced Protective Heterosubtypic Immunity From Cellular to Humoral Cross-Reactive Immune Responses. Front Immunol. 2020;11:1166.

24. Chang LA, Choi A, Rathnasinghe R, et al. Influenza breakthrough infection in vaccinated mice is characterized by non-pathological lung eosinophilia. Front Immunol. 2023;14:1217181.

25. Mesnil C, Raulier S, Paulissen G, et al. Lung-resident eosinophils represent a distinct regulatory eosinophil subset. J Clin Invest. 2016;126:3279–3295.

26. Lei C, Yang J, Hu J, et al. On the calculation of TCID50 for quantitation of virus infectivity. Virol Sin. 2021;36:141– 144.

27. Prophet EB, Bob, Arrington JB, et al. American Registry of Pathology. Laboratory Methods in Histotechnology.

28. Yeung ST, Ovando LJ, Russo AJ, et al. CD169+ macrophage intrinsic IL-10 production regulates immune homeostasis during sepsis. Cell Rep. 2023;42:112171.

29. Premeaux TA, Yeung ST, Pillai SK, et al. Elevated Galectin-9 across the human brain correlates with HIV neuropathology and detrimental cognitive states. J Neurovirol. 2023;29:337–345.

30. Kanda A, Yun Y, Van Bui D, et al. The multiple functions and subpopulations of eosinophils in tissues under steady-state and pathological conditions. Allergol Int. 2021;70:9–18.

31. Shah K, Ignacio A, McCoy KD, et al. The emerging roles of eosinophils in mucosal homeostasis. Mucosal Immunol. 2020;13:574–583.

32. Giacalone VD, Margaroli C, Mall MA, et al. Neutrophil adaptations upon recruitment to the lung: New concepts and implications for homeostasis and disease. Int J Mol Sci. 2020;21:851.

33. Johansson C, Kirsebom FCM. Neutrophils in respiratory viral infections. Mucosal Immunol. 2021;14:815–827.

34. Draijer C, Peters-Golden M. Alveolar macrophages in allergic asthma: The forgotten cell awakes. Curr Allergy Asthma Rep. 2017;17:12.

35. Panahipoor Javaherdehi A, Ghanbari S, Mahdavi P, et al. The role of alveolar macrophages in viral respiratory infections and their therapeutic implications. Biochem Biophys Rep. 2024;40:101826.

36. Feo-Lucas L, Godio C, Minguito de la Escalera M, et al. Airway allergy causes alveolar macrophage death, profound alveolar disorganization and surfactant dysfunction. Front Immunol. 2023;14:1125984.

37. Ghoneim HE, Thomas PG, McCullers JA. Depletion of alveolar macrophages during influenza infection facilitates bacterial superinfections. J Immunol. 2013;191:1250–1259.

38. Hashimoto D, Chow A, Noizat C, et al. Tissue-resident macrophages self-maintain locally throughout adult life with minimal contribution from circulating monocytes. Immunity. 2013;38:792–804.

39. David C, Verney C, Si-Tahar M, et al. The deadly dance of alveolar macrophages and influenza virus. Eur Respir Rev;33. Epub ahead of print October 2024. DOI: 10.1183/16000617.0132-2024.

40. Califano D, Furuya Y, Metzger DW. Effects of influenza on alveolar macrophage viability are dependent on mouse genetic strain. J Immunol. 2018;201:134–144.

41. Gurram RK, Zhu J. Orchestration between ILC2s and Th2 cells in shaping type 2 immune responses. Cell Mol Immunol. 2019;16:225–235.

42. Lloyd CM, Snelgrove RJ. Type 2 immunity: Expanding our view. Sci Immunol. 2018;3:eaat1604.

43. Chetty A, Darby MG, Pillaye J, et al. Induction of Siglec-FhiCD101hi eosinophils in the lungs following murine hookworm Nippostrongylus brasiliensis infection. Front Immunol. 2023;14:1170807.

44. Ganga L, Sharma P, Tiwari S, et al. Immunophenotypic and functional characterization of eosinophil and migratory dendritic cell subsets during filarial manifestation of tropical pulmonary eosinophilia. ACS Infect Dis. 2023;9:1105–1122.

45. Noble S-L, Vacca F, Hilligan KL, et al. Helminth infection induces a distinct subset of CD101hi lung tissue-infiltrating eosinophils that are differentially regulated by type 2 cytokines. Immunol Cell Biol. 2024;102:734–746.

46. Gupta RK. Aluminum compounds as vaccine adjuvants. Adv Drug Deliv Rev. 1998;32:155–172.

47. Snapper CM, Paul WE. Interferon-gamma and B cell stimulatory factor-1 reciprocally regulate Ig isotype production. Science. 1987;236:944–947.

48. Germann T, Gately MK, Schoenhaut DS, et al. Interleukin-12/T cell stimulating factor, a cytokine with multiple effects on T helper type 1 (Th1) but not on Th2 cells. Eur J Immunol. 1993;23:1762–1770.

49. Stevens TL, Bossie A, Sanders VM, et al. Regulation of antibody isotype secretion by subsets of antigen-specific helper T cells. Nature. 1988;334:255–258.

50. Hoeksema M, Tripathi S, White M, et al. Arginine-rich histones have strong antiviral activity for influenza A viruses. Innate Immun. 2015;21:736–745.

51. Zafarani A, Razizadeh MH, Haghi A. Neutrophil extracellular traps in influenza infection. Heliyon. 2023;9:e23306.

52. Ashar HK, Mueller NC, Rudd JM, et al. The role of extracellular histones in influenza virus pathogenesis. Am J Pathol. 2018;188:135–148.

53. Delgado-Rizo V, Martínez-Guzmán MA, Iñiguez-Gutierrez L, et al. Neutrophil extracellular traps and its implications in inflammation: An overview. Front Immunol. 2017;8:81.

54. Yousefi S, Simon D, Simon H-U. Eosinophil extracellular DNA traps: molecular mechanisms and potential roles in disease. Curr Opin Immunol. 2012;24:736–739.

55. Mukherjee M, Lacy P, Ueki S. Eosinophil extracellular traps and inflammatory pathologies-untangling the web! Front Immunol. 2018;9:2763.

56. Shen K, Zhang M, Zhao R, et al. Eosinophil extracellular traps in asthma: implications for pathogenesis and therapy. Respir Res. 2023;24:231.

57. Arnold IC, Artola-Borán M, Tallón de Lara P, et al. Eosinophils suppress Th1 responses and restrict bacterially induced gastrointestinal inflammation. J Exp Med. 2018;215:2055–2072.

58. Grilz E, Mauracher L-M, Posch F, et al. Citrullinated histone H3, a biomarker for neutrophil extracellular trap formation, predicts the risk of mortality in patients with cancer. Br J Haematol. 2019;186:311–320.

59. Rathnasinghe R, Jangra S, Ye C, et al. Characterization of SARS-CoV-2 Spike mutations important for infection of mice and escape from human immune sera. Nat Commun. 2022;13:3921.

60. Annunziato F, Romagnani C, Romagnani S. The 3 major types of innate and adaptive cell-mediated effector immunity. J Allergy Clin Immunol. 2015;135:626–635.

61. Wiese AV, Duhn J, Korkmaz RÜ, et al. C5aR1 activation in mice controls inflammatory eosinophil recruitment and functions in allergic asthma. Allergy. 2023;78:1893–1908.

62. Noble S-L, Mules TC, Le Gros G, et al. The immunoregulatory potential of eosinophil subsets. Immunol Cell Biol. 2024;102:775–786.

63. Ruegg CL, Rivas A, Madani ND, et al. V7, a novel leukocyte surface protein that participates in T cell activation. II. Molecular cloning and characterization of the V7 gene. J Immunol. 1995;154:4434–4443.

64. Bouloc A, Bagot M, Delaire S, et al. Triggering CD101 molecule on human cutaneous dendritic cells inhibits T cell proliferation via IL-10 production. Eur J Immunol. 2000;30:3132–3139.

65. Schey R, Dornhoff H, Baier JLC, et al. CD101 inhibits the expansion of colitogenic T cells. Mucosal Immunol. 2016;9:1205–1217.

66. Conroy DM, Williams TJ. Eotaxin and the attraction of eosinophils to the asthmatic lung. Respir Res. 2001;2:150– 156.

67. Wang Q, Liu J. Regulation and immune function of IL-27. Adv Exp Med Biol. 2016;941:191–211.

68. Jafarzadeh A, Nemati M, Jafarzadeh S, et al. The immunomodulatory potentials of interleukin-27 in airway allergies. Scand J Immunol. 2021;93:e12959.

69. Amsden H, Kourko O, Roth M, et al. Antiviral activities of interleukin-27: A partner for interferons? Front Immunol. 2022;13:902853.

70. Rangel-Moreno J, Hartson L, Navarro C, et al. Inducible bronchus-associated lymphoid tissue (iBALT) in patients with pulmonary complications of rheumatoid arthritis. J Clin Invest. 2006;116:3183–3194.

71. Cayrol C, Girard J-P. Interleukin-33 (IL-33): A critical review of its biology and the mechanisms involved in its release as a potent extracellular cytokine. Cytokine. 2022;156:155891.

72. Le Goffic R, Arshad MI, Rauch M, et al. Infection with influenza virus induces IL-33 in murine lungs. Am J Respir Cell Mol Biol. 2011;45:1125–1132.

73. Bunting MM, Shadie AM, Flesher RP, et al. Interleukin-33 drives activation of alveolar macrophages and airway inflammation in a mouse model of acute exacerbation of chronic asthma. Biomed Res Int. 2013;2013:250938.

74. De Giovanni M, Dang EV, Chen KY, et al. Platelets and mast cells promote pathogenic eosinophil recruitment during invasive fungal infection via the 5-HIAA-GPR35 ligand-receptor system. Immunity. 2023;56:1548–1560.e5.

75. Rothenberg ME, Hogan SP. The eosinophil. Annu Rev Immunol. 2006;24:147–174.

76. Scott G, Asrat S, Allinne J, et al. IL-4 and IL-13, not eosinophils, drive type 2 airway inflammation, remodeling and lung function decline. Cytokine. 2023;162:156091.

77. Bowen JL, Keck K, Baruah S, et al. Eosinophil expression of triggering receptor expressed on myeloid cells 1 (TREM-1) restricts type 2 lung inflammation. J Leukoc Biol. 2024;116:409–423.

