## Supplementary Figures for "Transient lung eosinophilia during breakthrough influenza infection in vaccinated mice is associated with protective and balanced Type 1/2 immune responses"

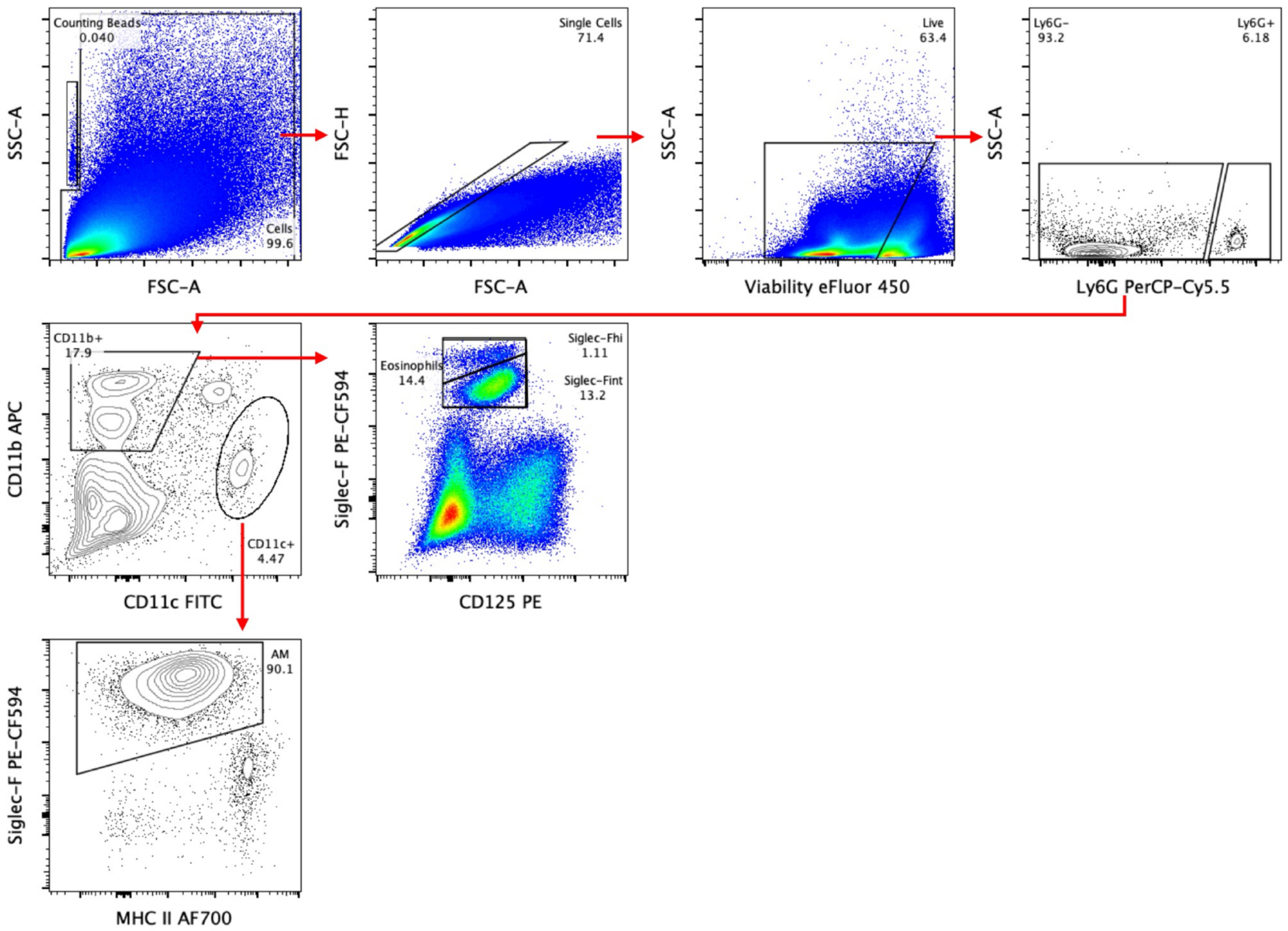


**Figure S1.** Gating strategy for flow cytometry of the lungs. Representative mouse from the breakthrough infection group at 7 DPC is shown here.

**A**

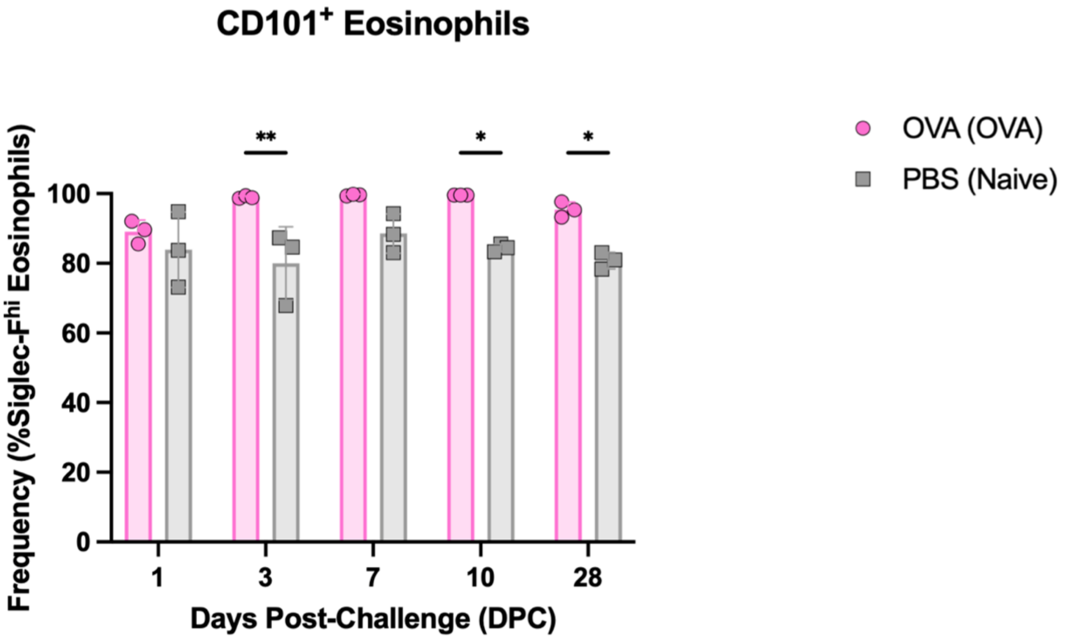


**B**

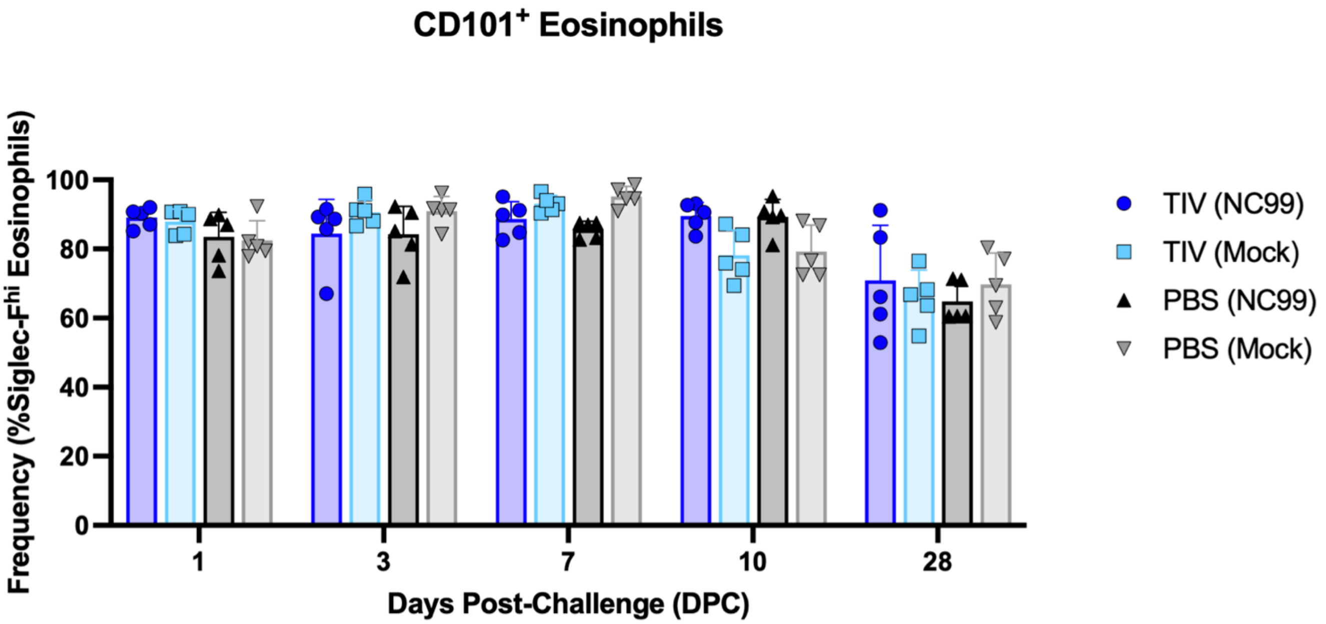


**Figure S2. The majority of Siglec-F$^{hi}$ eosinophils express CD101.** Frequency of CD101$^+$ cells within the Siglec-F$^{hi}$ eosinophil population was measured for (**A**) OVA sensitized mice or (**B**) breakthrough infection mice and controls. Statistical significance was determined via ordinary two-way ANOVA (**A**) with Šídák's multiple comparisons test with a single pooled variance or (**B**) with Tukey's multiple comparisons test with a single pooled variance. **P = 0.001 to 0.01, *P = 0.01 to 0.05.

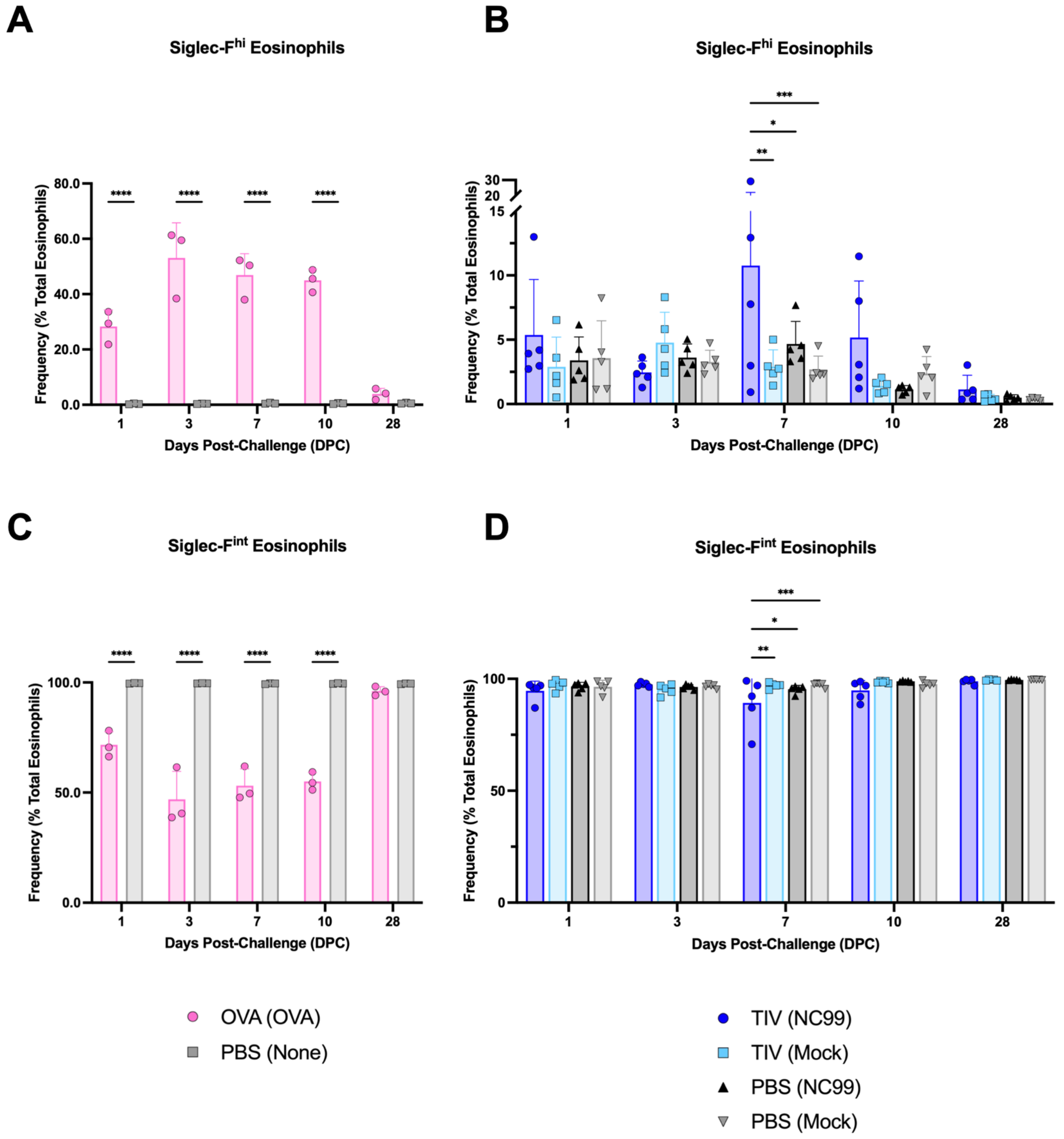


**Figure S3. Significant enrichment for the Siglec-F$^{hi}$ subset of eosinophils only occurs in OVA sensitized mice or breakthrough infection mice.** Frequency of Siglec-F$^{hi}$ eosinophils within the total eosinophil population in (**A**) OVA-sensitized mice and controls or (**B**) breakthrough infection mice and controls. Frequency of Siglec-F$^{hi}$ eosinophils within the total eosinophil population in (**C**) OVA-sensitized mice and controls or (**D**) breakthrough infection mice and controls. Statistical significance was determined via ordinary two-way ANOVA (**A**, **C**) with Šídák's multiple comparisons test with a single pooled variance or (**B, D**) with Tukey's multiple comparisons test with a single pooled variance. ****P < 0.0001, ***P = 0.0001 to 0.001, **P = 0.001 to 0.01, *P = 0.01 to 0.05.

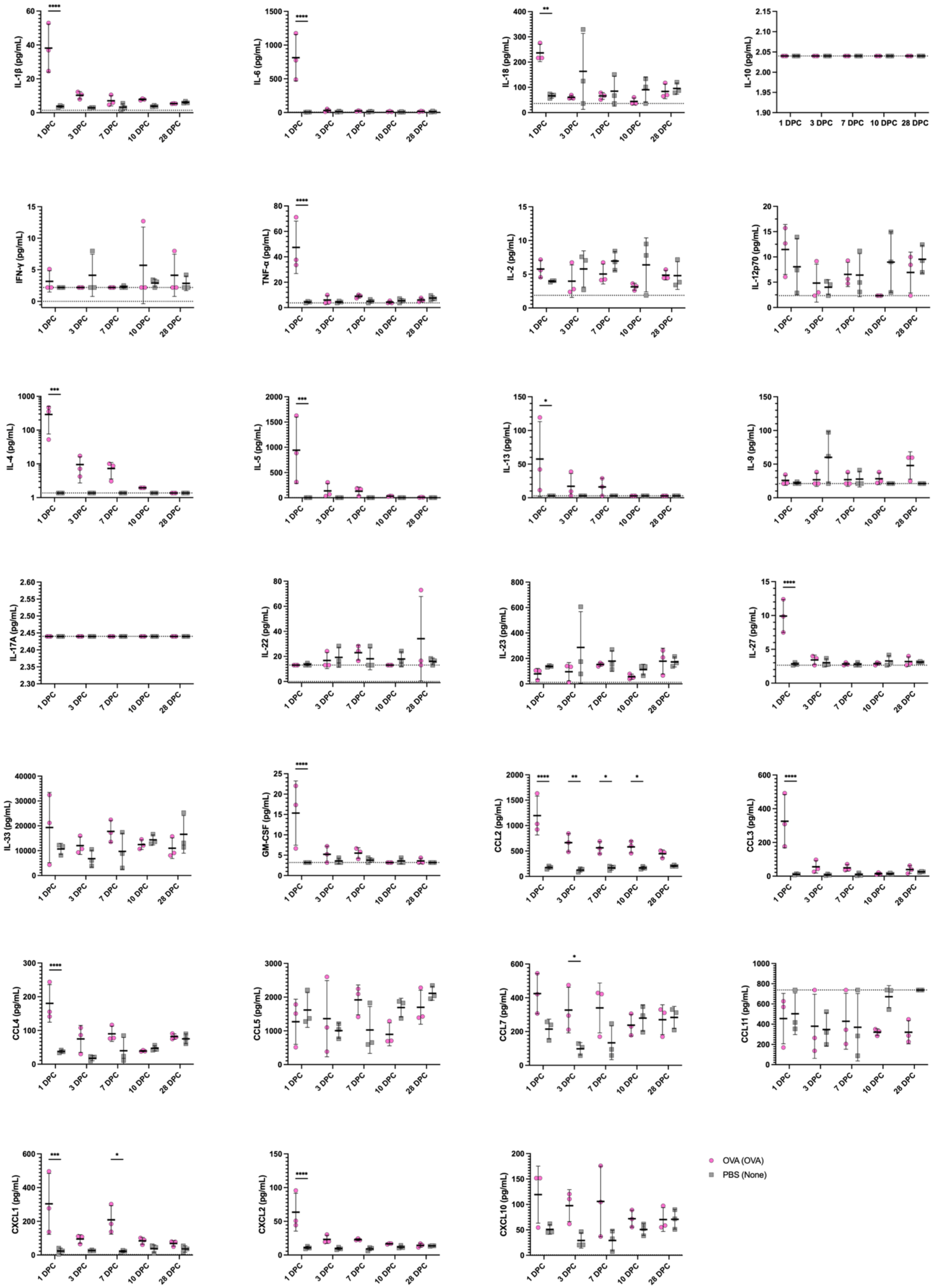
IL-1β (pg/mL)
IL-6 (pg/mL)
IL-18 (pg/mL)
IL-10 (pg/mL)
IFN-γ (pg/mL)
TNF-α (pg/mL)
IL-2 (pg/mL)
IL-12p70 (pg/mL)
IL-4 (pg/mL)
IL-5 (pg/mL)
IL-13 (pg/mL)
IL-9 (pg/mL)
IL-17A (pg/mL)
IL-22 (pg/mL)
IL-23 (pg/mL)
IL-27 (pg/mL)
IL-33 (pg/mL)
GM-CSF (pg/mL)
CCL2 (pg/mL)
CCL3 (pg/mL)
CCL4 (pg/mL)
CCL5 (pg/mL)
CCL7 (pg/mL)
CCL11 (pg/mL)
CXCL1 (pg/mL)
CXCL2 (pg/mL)
CXCL10 (pg/mL)
1 DPC
3 DPC
7 DPC
10 DPC
28 DPC
OVA (OVA)
PBS (None)

**Figure S4. Cytokine and chemokine concentrations in lung homogenate supernatants from OVA-sensitized mice and controls.** Statistical significance was determined via ordinary two-way ANOVA with Šídák's multiple comparisons test with a single pooled variance. ****$P < 0.0001$, ***$P = 0.0001$ to 0.001, **$P = 0.001$ to 0.01, *$P = 0.01$ to 0.05.

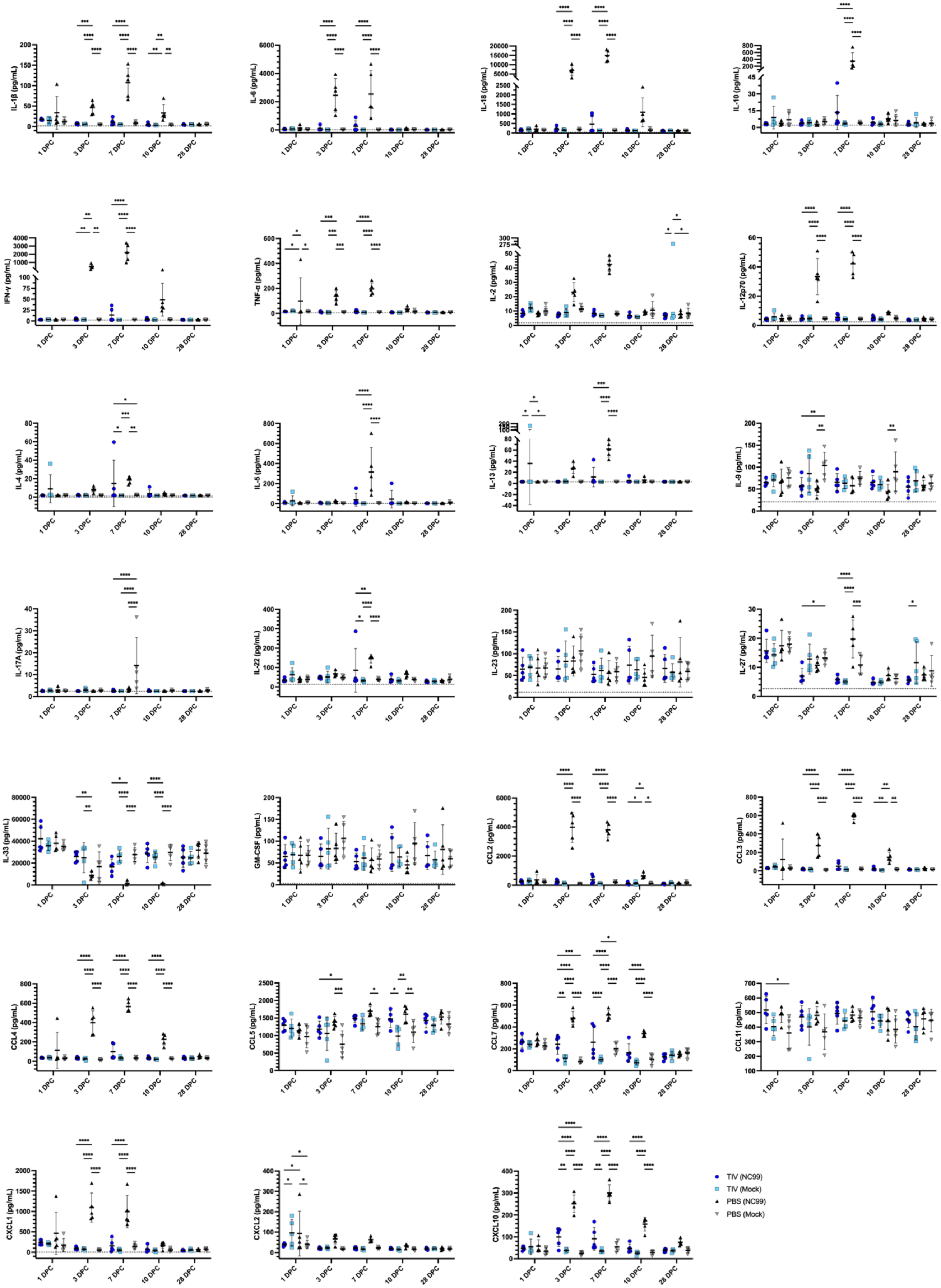

IL-1β (pg/mL)
IL-6 (pg/mL)
IL-18 (pg/mL)
IL-10 (pg/mL)
IFN-γ (pg/mL)
TNF-α (pg/mL)
IL-2 (pg/mL)
IL-12p70 (pg/mL)
IL-4 (pg/mL)
IL-5 (pg/mL)
IL-13 (pg/mL)
IL-9 (pg/mL)
IL-17A (pg/mL)
IL-22 (pg/mL)
IL-23 (pg/mL)
IL-27 (pg/mL)
IL-33 (pg/mL)
GM-CSF (pg/mL)
CCL2 (pg/mL)
CCL3 (pg/mL)
CCL4 (pg/mL)
CCL5 (pg/mL)
CCL7 (pg/mL)
CCL11 (pg/mL)
CXCL1 (pg/mL)
CXCL2 (pg/mL)
CXCL10 (pg/mL)
1 DPC
3 DPC
7 DPC
10 DPC
28 DPC
TIV (NC99)
TIV (Mock)
PBS (NC99)
PBS (Mock)

**Figure S5. Cytokine and chemokine concentrations in lung homogenate supernatants from breakthrough infection mice and controls.** Statistical significance was determined via ordinary two-way ANOVA with Tukey's multiple comparisons test with a single pooled variance. ****P < 0.0001, ***P = 0.0001 to 0.001, **P = 0.001 to 0.01, *P = 0.01 to 0.05.

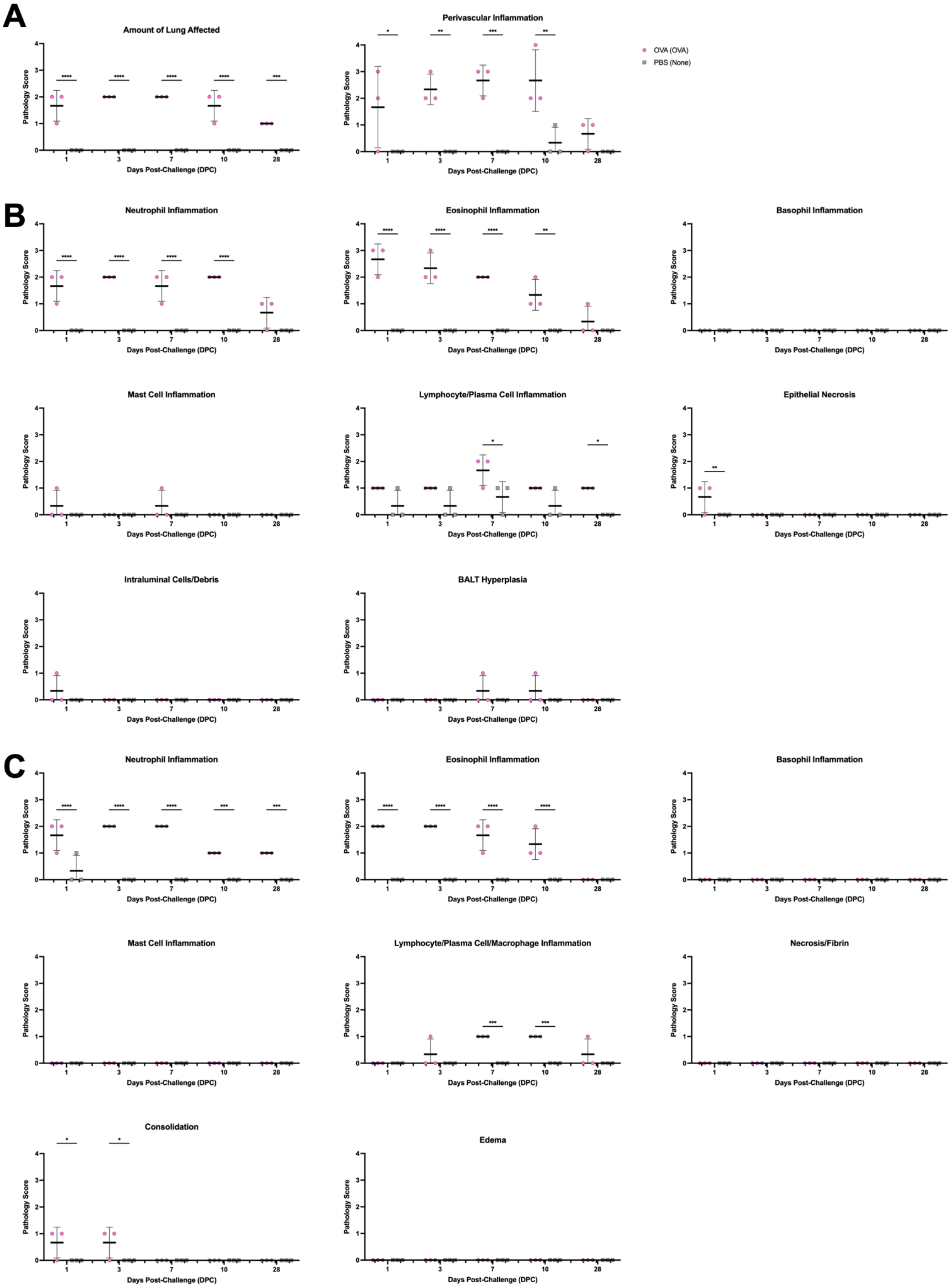

A
Amount of Lung Affected
Perivascular Inflammation
OVA (OVA)
PBS (None)
B
Neutrophil Inflammation
Eosinophil Inflammation
Basophil Inflammation
Mast Cell Inflammation
Lymphocyte/Plasma Cell Inflammation
Epithelial Necrosis
Intraluminal Cells/Debris
BALT Hyperplasia
C
Neutrophil Inflammation
Eosinophil Inflammation
Basophil Inflammation
Mast Cell Inflammation
Lymphocyte/Plasma Cell/Macrophage Inflammation
Necrosis/Fibrin
Consolidation
Edema
Pathology Score
Days Post-Challenge (DPC)

**Figure S6. Pathology scores for OVA-sensitized mice and controls.** Scores for metrics for the (**A**) total lung; (**B**) bronchi and bronchioles, peribronchial and peribronchiolar regions; (**C**) alveoli and alveolar septa. Statistical significance was determined via ordinary two-way ANOVA with Šídák's multiple comparisons test with a single pooled variance. ****P < 0.0001, ***P = 0.0001 to 0.001, **P = 0.001 to 0.01, *P = 0.01 to 0.05.

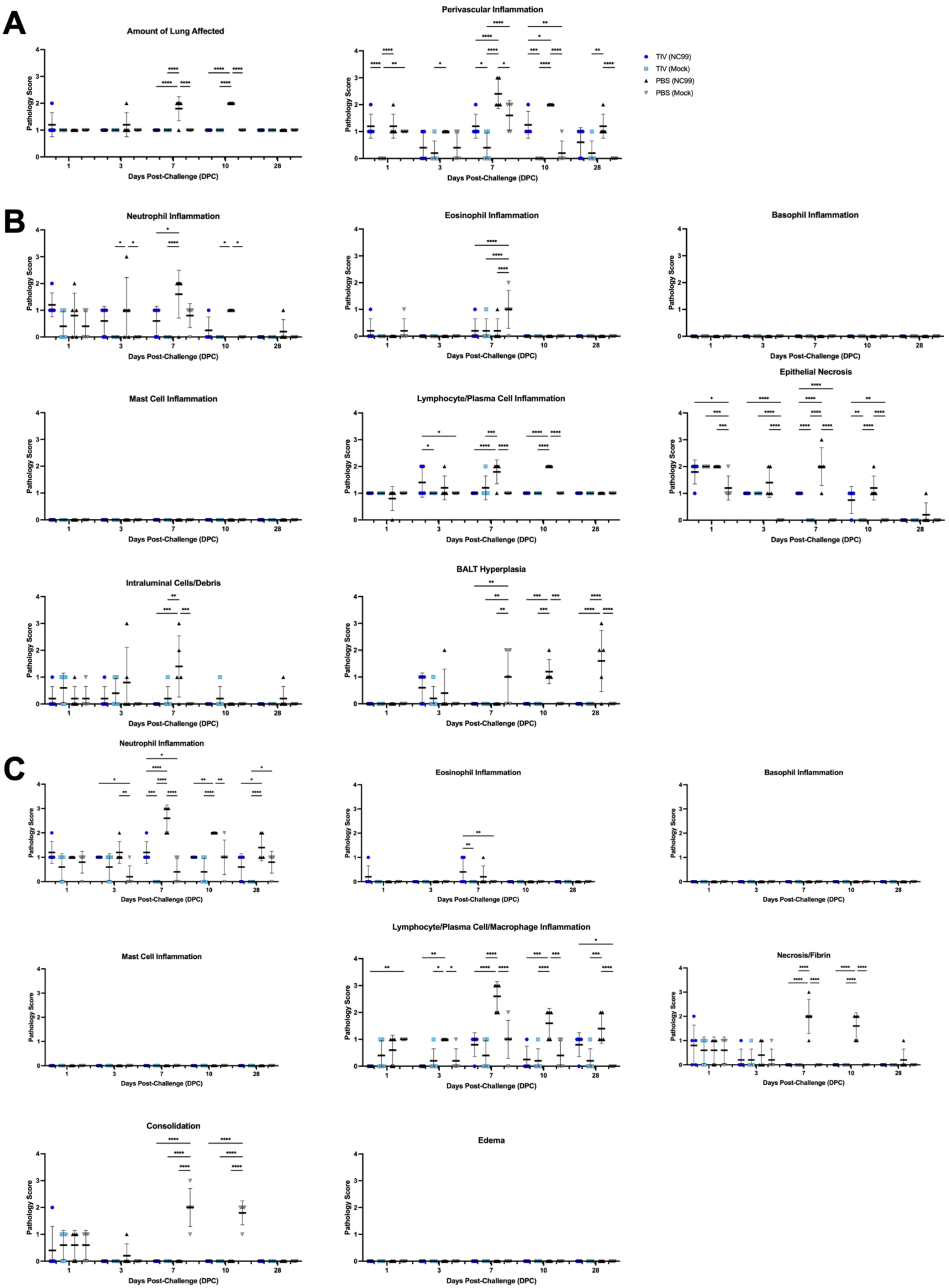

A
Amount of Lung Affected
Perivascular Inflammation
TIV (NC99)
TIV (Mock)
PBS (NC99)
PBS (Mock)
B
Neutrophil Inflammation
Eosinophil Inflammation
Basophil Inflammation
Mast Cell Inflammation
Lymphocyte/Plasma Cell Inflammation
Epithelial Necrosis
Intraluminal Cells/Debris
BALT Hyperplasia
C
Neutrophil Inflammation
Eosinophil Inflammation
Basophil Inflammation
Mast Cell Inflammation
Lymphocyte/Plasma Cell/Macrophage Inflammation
Necrosis/Fibrin
Consolidation
Edema
Pathology Score
Days Post-Challenge (DPC)

**Figure S7. Pathology scores for breakthrough infection mice and controls.** Scores for metrics for the (**A**) total lung; (**B**) bronchi and bronchioles, peribronchial and peribronchiolar regions; (**C**) alveoli and alveolar septa. Statistical significance was determined via ordinary two-way ANOVA with Tukey's multiple comparisons test with a single pooled variance. ****P < 0.0001, ***P = 0.0001 to 0.001, **P = 0.001 to 0.01, *P = 0.01 to 0.05.

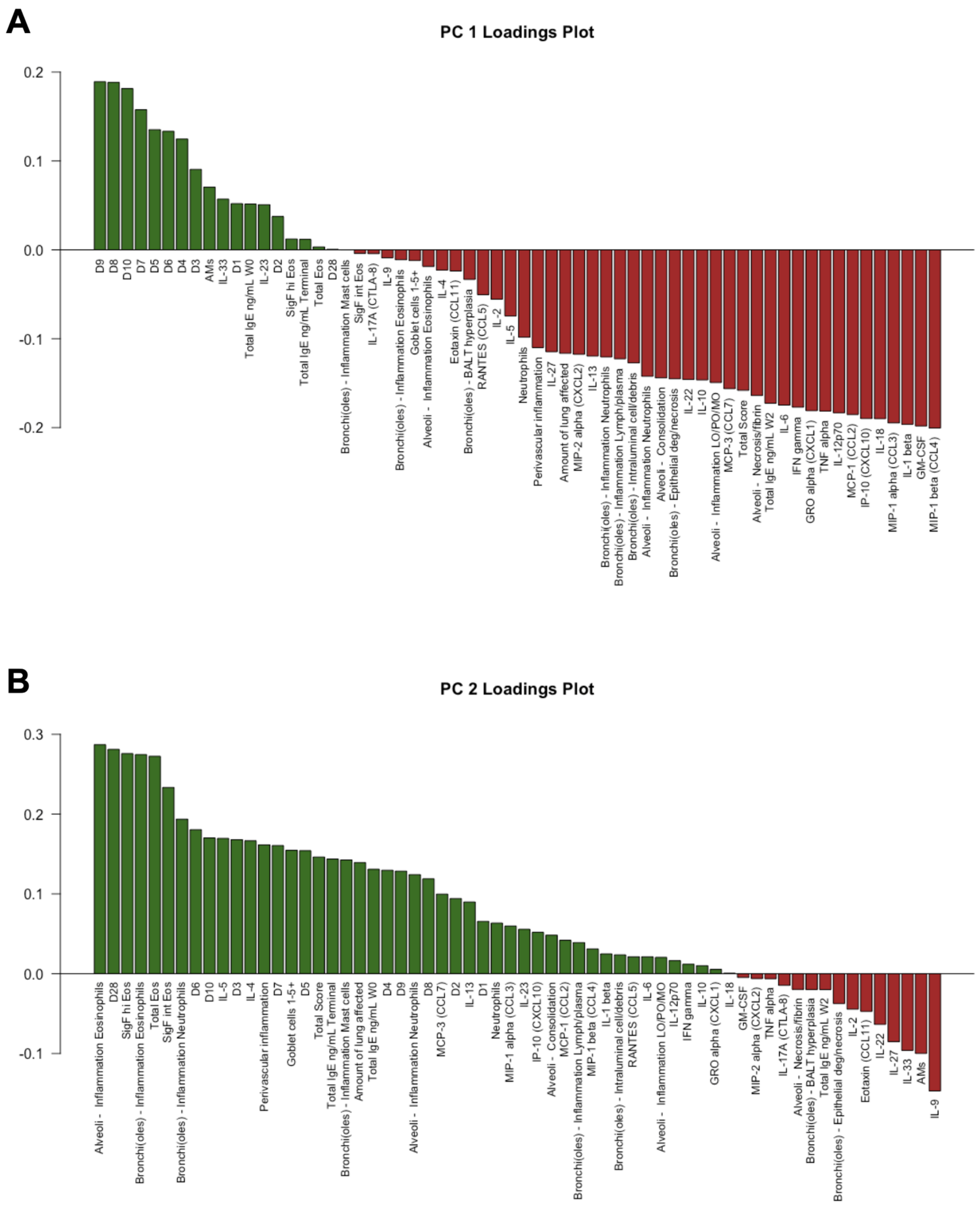


**Figure S8. PCA loadings.** Bar plot of loadings for (**A**) PC1 and (**B**) PC2.

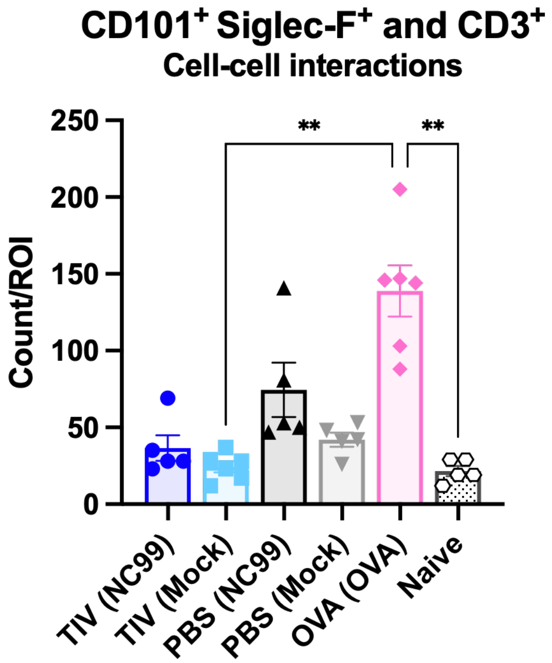


**Figure S9. CD101$^+$Siglec-F$^+$ and CD3$^+$ cell-cell interactions quantified per region of interest.** Statistical significance was determined using a Kruskal-Wallis test with Dunn's multiple comparisons test. **P = 0.001 to 0.01.
